# A user manual to measure gas diffusion kinetics in plants: Pneumatron construction, operation and data analysis

**DOI:** 10.1101/2021.02.08.430283

**Authors:** C.L. Trabi, L. Pereira, X. Guan, M.T. Miranda, P.R.L. Bittencourt, R.S. Oliveira, R.V. Ribeiro, S. Jansen

## Abstract

The Pneumatron device presented measures gas diffusion kinetics in the xylem of plants. The device provides an easy, low-cost, and powerful tool for research on plant water relations. Here, we describe in detail how to construct and operate this device to estimate xylem vulnerability to embolism, and how to analyse pneumatic data. Simple and more elaborated ways of constructing a Pneumatron are shown, either using wires, a breadboard, or a printed circuit board. The instrument is based on an open-source hardware and software system, which allows users to operate it in an automated or semi-automated way. A step-by-step manual and a troubleshooting section are provided. An excel spreadsheet and an R-script are also presented for fast and easy data analysis. This manual should help new users to avoid common mistakes, especially regarding stable measurements of the minimum and maximum amount of gas that can be discharged from xylem tissue.

## 1 Introduction

The Pneumatron is a device that allows automated measurements of the gas diffusion kinetics in plant xylem tissue (Pereira et al., 2020a). Pneumatic measurements have been applied to xylem tissue of various plant organs, such as stems, roots, and leaves, to estimate vulnerability to hydraulic failure of the water conducting cells, which is especially relevant to plants that undergo drought stress. Although the device has been designed to address questions in the field of xylem anatomy and physiology, it is expected that the instrument can also be applied to a wide range of other porous media.

During a pneumatic method measurement, a partial vacuum is pulled in a discharge tube to extract gas from xylem tissue of a cut branch, petiole, or root during less than one minute (Pereira et al., 2016; Bittencourt et al., 2018). The amount of gas extracted can be calculated from pressure measurements, using the ideal gas law. The Pneumatron has been shown to provide a major improvement of the manual pneumatic apparatus because the vacuum condition, pressure sensor readings, data storage, and the opening or closing of valves can be done within milliseconds. By applying repetitive measurements over time, and combining these pneumatic data with a quantification of sample dehydration, a “vulnerability curve” can be obtained in a straightforward way. For a general understanding of the manual pneumatic method, we refer to literature (Pereira et al., 2016, 2020a; Bittencourt et al., 2018; Zhang et al., 2018; Jansen et al., 2020).

So far, pneumatic vulnerability curves have been conducted for hundreds of species by few research groups. Although its construction is simple and based on an accessible and open-source platform (Arduino), there is a need for a detailed manual with clear instructions on the construction of a Pneumatron device, its operation, and the analysis of pneumatic data. Such details are crucial to introduce new research group to pneumatic measurements, which are different from measurements of hydraulic conductivity, and to ensure accurate and correct interpreation of the data obtained. This paper aims to provide such a user manual, and will avoid common mistakes in pneumatic experiments and misinterpretation of measurements. Besides its importance for measuring xylem embolism resistance, the Pneumatron can also be used to study gas kinetics of plants *in vivo*, and vessel length distributions (Pereira et al., 2020b; Yang et al., Submitted). These methods require minor modifications of the software programme and tube connections. Contrary to other standard methods on embolism resistance, the Pneumatron device is very fast and user-friendly, which makes this device also useful for field measurements.

## 2 Materials and Equipment

### 2.1 Equipment

The Pneumatron is composed of a microcontroller system (Atmega328P, Microchip, on anArduino^®^ Uno board), a data storage (SD card) and real time clock (DS1307, Maxim Integrated; both assembled on an Adafruit^®^ Data Logger Shield), a 16 bits analog-to-digial converter (ADS1115, Texas Instrument), a vacuum pump, a solenoid valve (3/2 connection normal closed) and its driver (logic-level N-channel mosfet plus flyback diode) (Fig. 1a). These components are easily available in electronic shops.

**Figure 1:**
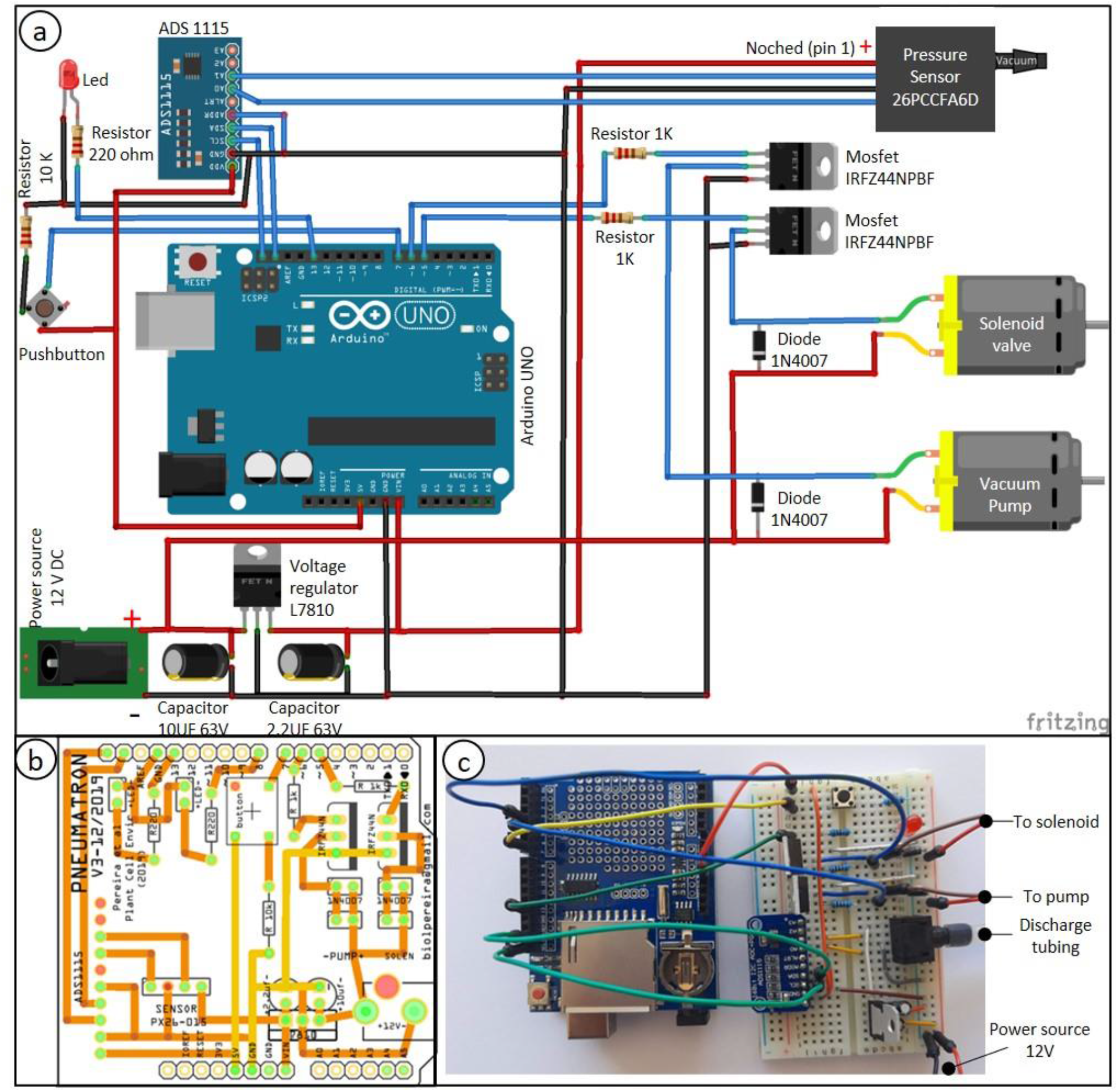
Connection scheme between the Pneumatron components based on wires (a; the data logger module, which is connected on top of the Arduino board, is not shown), a two-layer printed circuit board (PCB) of a wireless Pneumatron shield (b) (https://github.com/Pneumatron/construction), and a non-permanent assembly using a breadboard (c). The complete list of all components is listed in Table S1 and a schematic connection among the parts is presented in Figure S1.

A printed circuit board (PCB) to construct the Pneumatron Shield (Fig. 1b), may be used to facilitate the connection among electronic parts, reducing connection problems and providing better electronic properties while requiring minimal soldering skills. The PCB can be easily manufactured via several PCB producers by using the production files (Gerber Files; available in https://github.com/Pneumatron/construction). Alternatively, it is possible to use a breadboard to make all connections (Fig.1c), although this is not recommended, as it increases the chance of failure due to bad contacts and, if not properly done, can lead to voltage gradients in the analog-to-digital converter and sensor supply and grounding. A breadboard is not a permanent option, but can be useful for testing or for a low demand experiment.

The individual components of the Pneumatron are listed in Table S1. The assembly and testing do not require specialized knowledge, but correct orientation of some components is must be respected (Fig 1a). For protecting the sensitive parts of the Pneumatron, it is advised to enclose these into a box. It is recommended to do this for preventing any damage to the vacuum pump and solenoid valve and potential short circuit.

After assembling the electronic parts (Fig. 1), the pressure sensor, the vacuum pump, and the solenoid valve need to be connected to each other with silicone tubing (Fig. 2a). These components are then connected to the PCB circuit board (Fig. 2a). Once you have assembled the pins onto the data logger shield and installed the different components onto the PCB board, you need to assemble them into a stacked-up structure with the Arduino at the bottom, the data logger shield in the middle, and the Pneumatron shield at the top (Fig. 2b).

**Figure 2:**
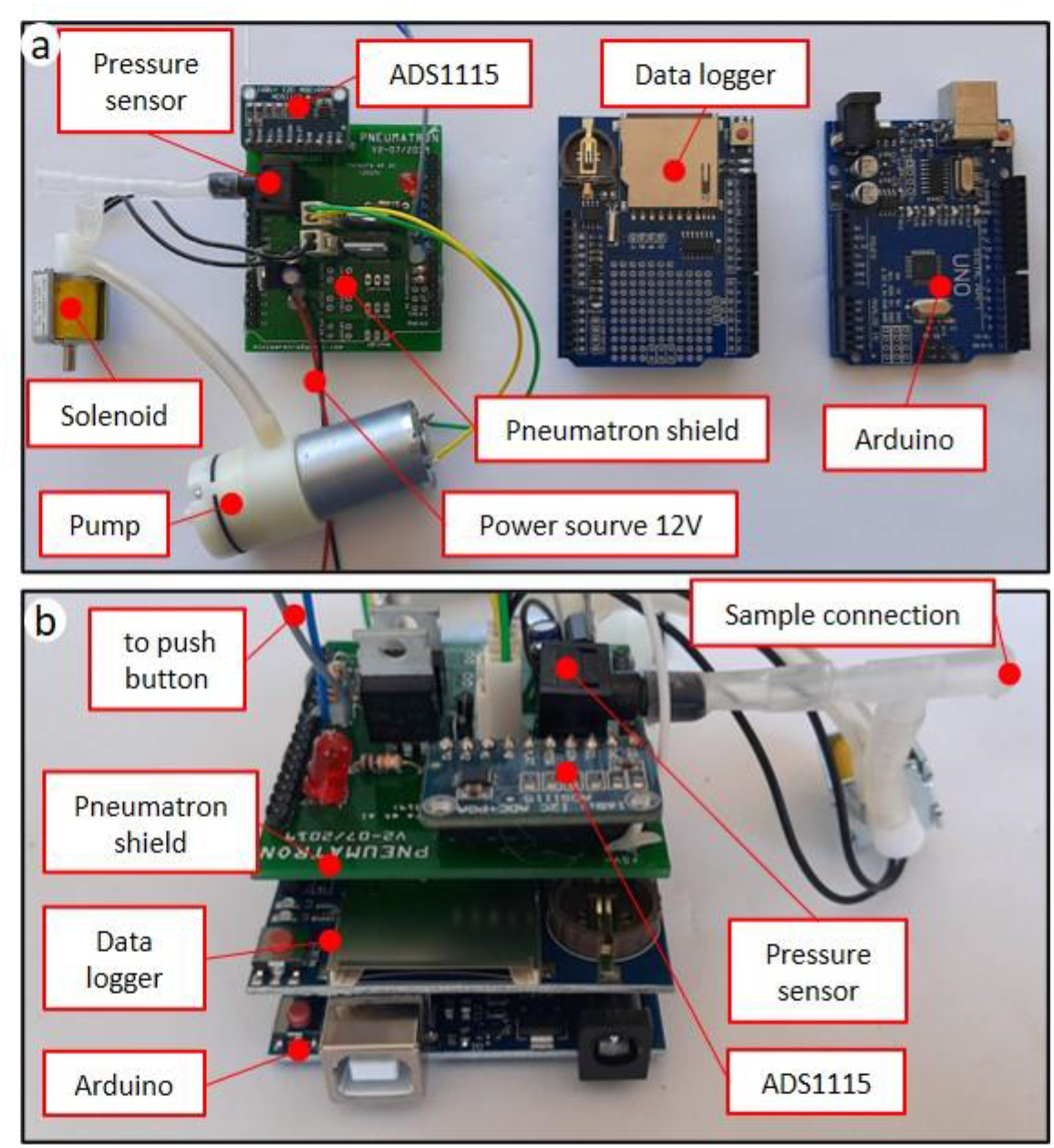
Components and connections (a) and assembling position (b) of the Pneumatron with a Pneumatron shield (in which the push-button and power source were connected using wires).

### 2.2 Software

The Pneumatron is built on an Arduino^®^ Uno or similar board. To be able to upload or modify the programmes, installation of the software is required. Further information can be found on the Arduino web-site (https://www.arduino.cc/en/Main/Software). System information and requirements are needed to get the correct version of the programme, depending on which platform the software is to be installed on.

In order to get the programme fully functioning, some of the libraries need to be installed or updated. The libraries used in this version of the Pneumatron are the SPI and and Wire (v. 2.3.5) for SPI and I2C communication, RTCLib (version 1.2.4) for using the DS1307 real-time clock, Adafruit_ADS1X15 (v. 1.0.1) for using the ADS1115 analog-to-digital converter and the SD library (v. 2.3.5) for using SD the card with FAT32 file system. More information can be found at: https://www.arduino.cc/en/guide/libraries. Arduino novices might require further training on how to upload a given programme onto the board. Some complementary information is available at: https://www.arduino.cc/en/main/howto.

The programmes for the Pneumatron can be downloaded from https://github.com/Pneumatron/software. Once you have uploaded the programme, your equipment is ready for calibration and testing.

### 2.3 Programmes

#### 2.3.1 Automated mode

The programme controls the pump and the solenoid valve, and reads data from the pressure sensor via an ADC 16 bits converter (Fig. 3). Firstly, the pump will be heard creating a partial vacuum inside the discharge tubing (ca. 40 kPa of absolute pressure), and should achieve this partial vacuum in less than one second. If not, either the vacuum pump is not working well, or there is a leakage. After reaching the target pressure, it will stop and the pressure values will be recorded for one minute (this period can be changed depending on the experimental requirements). During this measuring period, the LED will flash every 0.5 s. Finally, the apparatus pressure will return to atmospheric pressure via repeated opening and closing of the solenoid valve over a short time period (5 s), and the device will then wait for the next measurement to be taken (e.g., every 15 min). The suggested minimum time interval between measurements is 15 min, which allows for the gas in the xylem tissue to reequilibrate with atmospheric pressure before the next measurement.

**Figure 3:**
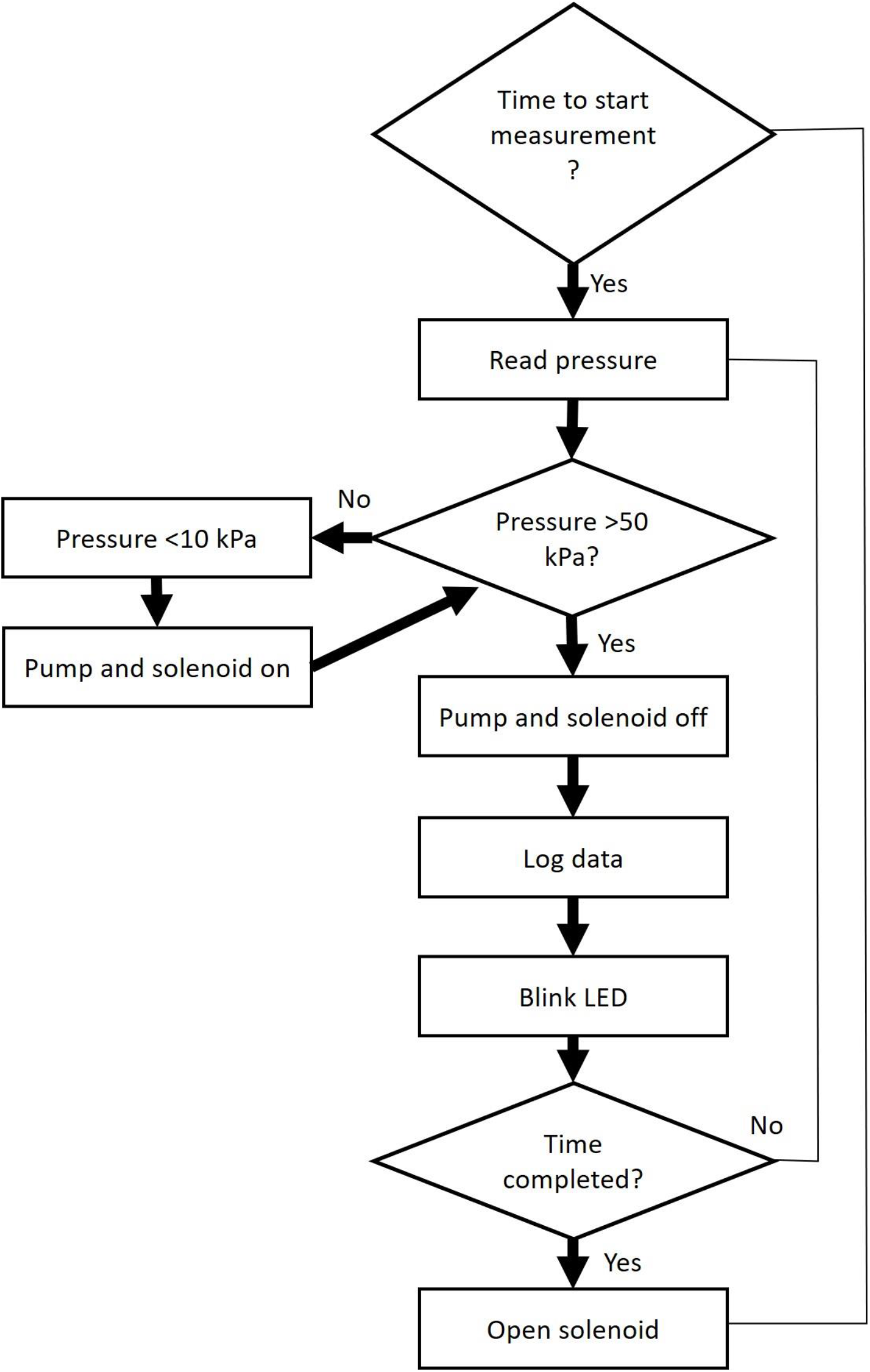
Workflow of a Pneumatron operation during automated mode.

#### 2.3.2 Semi-automated mode

The semi-automated mode is similar to the automated one, but each individual measurement has to be acttivated by the operator through a button. The rest of the semi-automated process is similiar to the automated method, except for activation of the solenoid valve at the end of the measurement. The button to take a measurement can only be pressed after an entire measuring cycle has been finished.

## 3 Methods

### 3.1 Testing and Calibration

#### 3.1.1 Testing the Pneumatron for leakage

After assembling the Pneumatron, the device should be tested for leakage. This is achieved by blocking off the tubing that is used for sample connection, while running the programme for several measurement cycles in the semi-automated mode. If the pump is working permanently, or frequently (e.g. more than once during a single measurement) please refer to the troubleshooting section (see the results section). Then, the device is powered off and the SD card removed. An ideal and fast way to test the data obtained is to import the csv file into excel and to select only the data column on the right, plotting the gas extraction data into a scatter point graph. The data should be as close to constant as possible (within 1 to 2%), apart from the first point, which is close to zero. If the data seems to be drifting more than 5% and decreasing regularly with each measurement, there is a leakage. More information about how to deal with leakages can be found in the troubleshooting section.

#### 3.1.2 Calibration

The Pneumatron measurements are based on relative values (percentage of air discharge) calculated from the difference between the initial and the final pressure measured in the discharge tubing. For this reason, the units used, that is volts or kPa, are not critical. However, if the correct pressure unit is required for a different purpose, it is necessary to calibrate the pressure sensor instead of using the manufacturer equation (default in the Pneumatron programmes) since the relation voltage/pressure changes depending on the voltage supply. To calibrate the sensor, use the calibration software (https://github.com/Pneumatron/software). There are two easy methods for calibration, one relying on the height of a water column (Fig. 4a), and the other one using a syringe pump (Fig. 4b).

**Figure 4:**
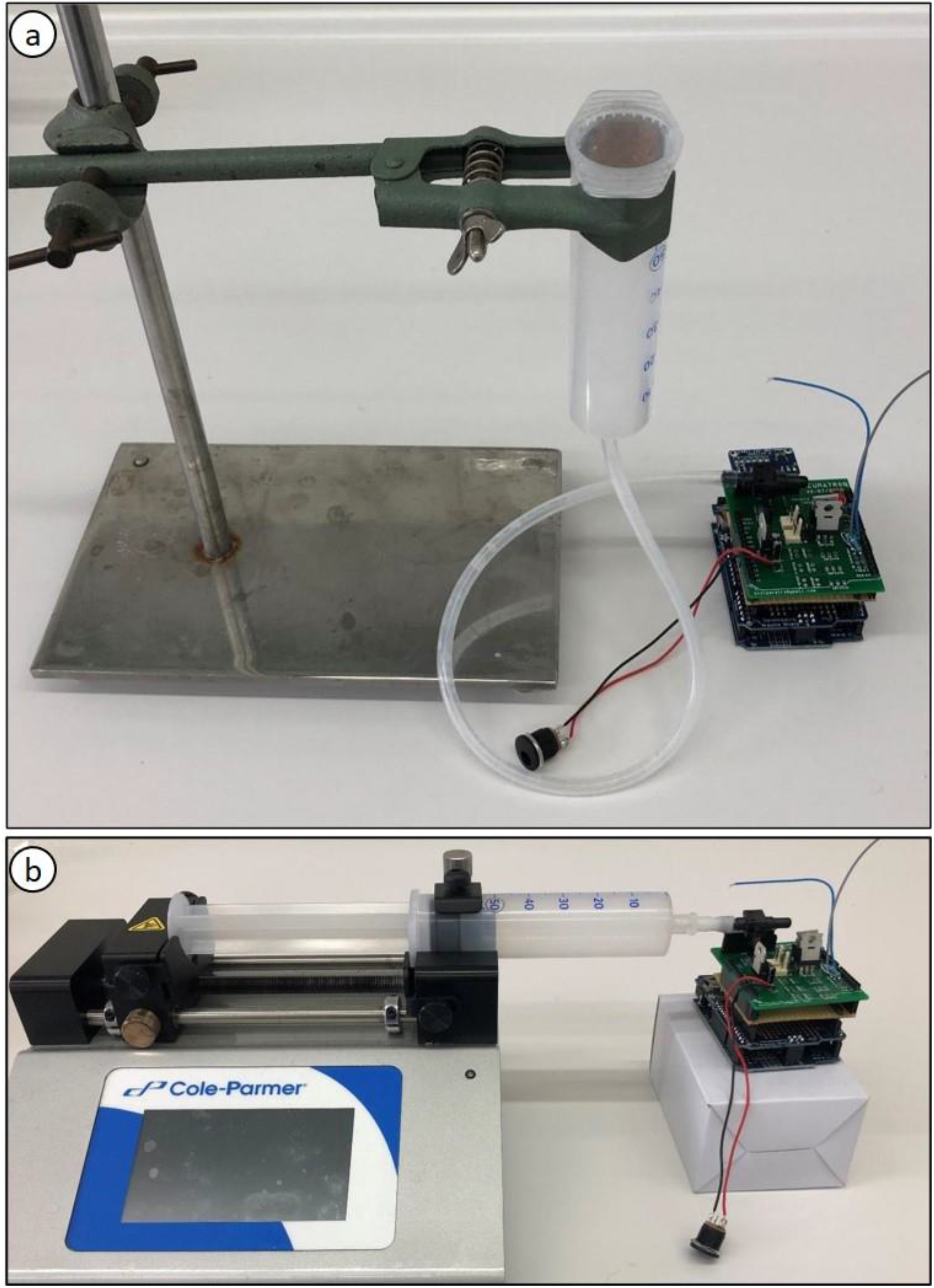
Calibration of the pressure sensor in a Pneumatron device using a water column (a), which requires any container with known height of the water column (such as a syringe without the plunger), tubing, a stand with a clamp to hold the container, and a ruler to measure the height of the water column (from the pressure sensor to the free surface of the liquid). Calibration of a pressure sensor can also be done using a syringe pump (b).

##### 3.1.2.1 Water column calibration

The pressure sensor (disconnected from the valve and pumping system, and reconnected to the tube with the syringe), the tubing, and the syringe should be fully filled with water. The syringe is then fixed by the stand at a certain height. The distance between the pressure sensor and the water surface of the liquid should then be measured. This process is repeated at two or three height levels.

When the measurements have been taken, the data should be extracted from the Pneumatron and the TXT file imported in any data plotter software. The pressure measurements should be represented by a step graph and the value for each step should be linked with the height of the water column. To determine the pressure that is produced by the water column, the following equation is used:

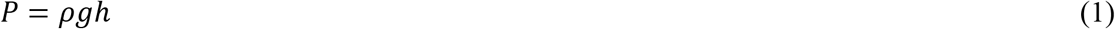

Where ρ is the density of the liquid used (1,000 kg m^-3^ for water), g is the acceleration of gravity (9.81 m s^-2^) and h is the height between the pressure sensor and the free surface of the liquid, in meter. The resulting pressure will be in Pa, and can be converted to kPa for ease of use (1,000 Pa = 1 kPa).

The calibration equation is obtained by determining fitting the applied pressure (y-axis) to the sensor voltage (x-axis). This equation (the slope and intercept values) should then be included in the programme (see instruction at the initial lines of the programme) and the programme uploaded once more onto the Arduino board.

##### 3.1.2.2 Syringe pump calibration

The Pneumatron is started and the air contained in the 60 mL syringe is compressed. The pressure measured is relative. When air in the 60 mL syringe is compressed to 50 mL, the pressure is then 20 kPa; at 40 mL, the pressure is 50 kPa, and 100 kPa is reached at 30 mL. The same process is followed as the water column calibration to determine the calibration equation and to upload the equation to the board.

##### 3.1.2.3 Implementation of the calibration parameters

Depending on the manufacturer of the pressure sensor, the sensor response for a given pressure will be different. In case the pressure sensor differs from the one suggested here, the datasheet should provide information about the response of the sensor to the pressure applied, as well as about the power supplied to the sensor, which is 10 volts in the Pneumatron. Please refer to the datasheet of the sensor and check the equation to be implemented in the programme (see https://github.com/Pneumatron/software), as shown in section 3.1.2.1.

### 3.2 Experimental Techniques

#### 3.2.1 Sample preparation

Unlike sampling for hydraulic measurements, plant material can be cut in air because the conduits that are cut open need to be filled with air. The cut open conduits therefore represent an extension of the discharge tube (Pereira et al., 2016; Jansen et al., 2020). The length of samples should be preferentially longer than the maximum vessel length (Greenidge, 1952), which is unproblematic if terminal organs such as terminal branches, roots, or single leaves are used as they are part of the end/start of the water transport system, and they are not showing another cut. Plant samples, however, should be intact and any damage due to natural or artificial wounds should be avoided. It is also possible to work with stem or root segments, as long as air-entry is prevented at one of the cut sides (Wu et al., 2020). The distal end of a segment could, for instance, be blocked with super glue or a resin to avoid gas exchange.

After cutting in air, samples should be covered up with a black plastic bag to avoid dehydration. Since it is important to start pneumatic measurements on samples that are well hydrated, samples need to be collected early in the morning, preferentially during the wet season. However, if these conditions are not possible, samples can be left within the plastic bag with their cut ends in water for several hours or one night before taking measurements.

#### 3.2.2 Determination of the discharge tubing volume

A crucial factor that determines the precision of the Pneumatron is the volume of the discharge tubing, which needs to be proportional to the amount of gas discharged from the xylem (Pereira et al., 2020a). To estimate the discharged tubing volume, you may use completely dehydrated organs, of similar size of the sample of interest, to measure the maximum amount of gas that can be discharged. This step is essential to prevent re-pumping when samples become considerably dehydrated, and to ensure maximun resolution of the pressure sensor (Pereira et al., 2020a). Repetitive measurements are required to make sure that the maximum amount of gas represents a constant plateau. The ideal volume of the discharge tube can then be estimated by dividing the maximum volume of gas discharged in microliters by 510.2 (Pereira et al., 2020a). It will give the volume in milliliters to maximize the Pneumatron precision.

Alternatively, you may take a few measurements on the same dehydrated plant organ using a small tubing volume (< 1 mL), and check if the vacuum pump will restart during the measurement time (e.g. within 1 min). If yes, you must increase the tubing volume and test it again. This step needs to be repeated until the pump is no longer reactivated within the 1 min interval. The volume can be adjusted by adding or removing a certain amount of rigid tubing. It is important that the tubing has a constant volume under the partial vacuum that is pulled. See Table S1 for tubing details. If the discharge tube is not adjusted, the measuring error during pneumatic experiments can be considerably high (Jansen et al., 2020; Pereira et al., 2021).

#### 3.2.3 Always test for leakage before you start

Close off the open tube of the Pneumatron, and insert an empty SD card to the data logger shield. Connect the Pneumatron to a power supply system. If the software uploaded follows the automated mode, you will immediately hear the vacuum pump. If not, you need to push the button to start the measurement. Then, wait for one min until the first measurement cycle has finished. The LED light flashes when pressure data are recorded, and stops when the measurement has finished. Then, remove the SD card and check the data to see if there was any potential leakage during the measurement (See details in how to get and analyse data, and the troubleshooting part for potential leakage).

#### 3.2.4 Cleaning the memory card

Because your data are always stored in the file “log.txt”, it is important to clean the memory card before you start with new measurements. Make sure that the SD card has been inserted properly (correct orientation and tight insertion).

#### 3.2.5 Connecting a sample to the Pneumatron

Using hydrated plant material, preferentially bagged to avoid rapid dehydration at the beginning, trim the cut end of the sample carefully with a fresh razor blade to have conduits nicely cut open. Cutting can be done in air as cut-open conduits should become embolised. When working with non-terminal branches, leaves, or roots, make sure that the length of the sample is longer than the maximum vessel length. Otherwise, a high amount of air will be sucked up via the cut-open conduit. As mentioned above, you also need to put glue at distal cuts or at broken twigs to avoid gas exchange via cut xylem. Altought it is possible that these cut or broken xylem structures are not connected to the cut-open xylem tissue to which the Pneumatron is connected, it is recommended to apply glue to these places.

Connect the branch end to the Pneumatron with elastic tubing (see Table S1 for details of the tubing and clamps), and choose the best-fitting tube, considering the size of your cut sample end. Use parafilm, plastic clamps, or glue to ensure that there is the smallest possible leakage.

Since dehydration of fresh samples is affected by environmental conditions, pneumatic measurement should ideally be done under conditions of stable temperature and humidity. As initial dehydration is typically fast, samples can be bagged (or semi-bagged). This is especially recommended when working with species that are rather vulnerable to embolism, and will allow you to collect more data points with the Pneumatron, especially at water potentials that are not very negative.

#### 3.2.6 Turning on the apparatus

Connect the apparatus to the power supply and the measurements will start automatically when following the automatic mode. Measurements will be taken at every 15 min by default, but the time interval can easily be changed in the programme. The calculation of the volume of air discharged for each measurement will be discussed below in the “Data analysis” section.

#### 3.2.7 Stopping the measurements

Measurements can be stopped when the branches are completely dehydrated (see the next section about water potential measurements). After severe dehydration, the maximum amount of gas extracted from the plant sample has been achieved, and there will be no longer increases in the percentage of gas discharged with further dehydration. Then, a curve between air discharge and time or, preferentially water potential, will form a plateau.

A common mistake is that measurements are stopped too early, before the maximum amount of extracted gas has been reached. In this case, the Ψ_50_ (water potential in which 50% of xylem embolism is formed) can be strongly underestimated as simulated in Figure 5. When working with a sample that has leaves, it is recommended to run the measurements until the leaves become crispy. At this dehydration point, it is also important to avoid leakages caused by stem shrinkage, tightening the clamp and reapplying glue at the connection with the Pneumatron.

**Figure 5:**
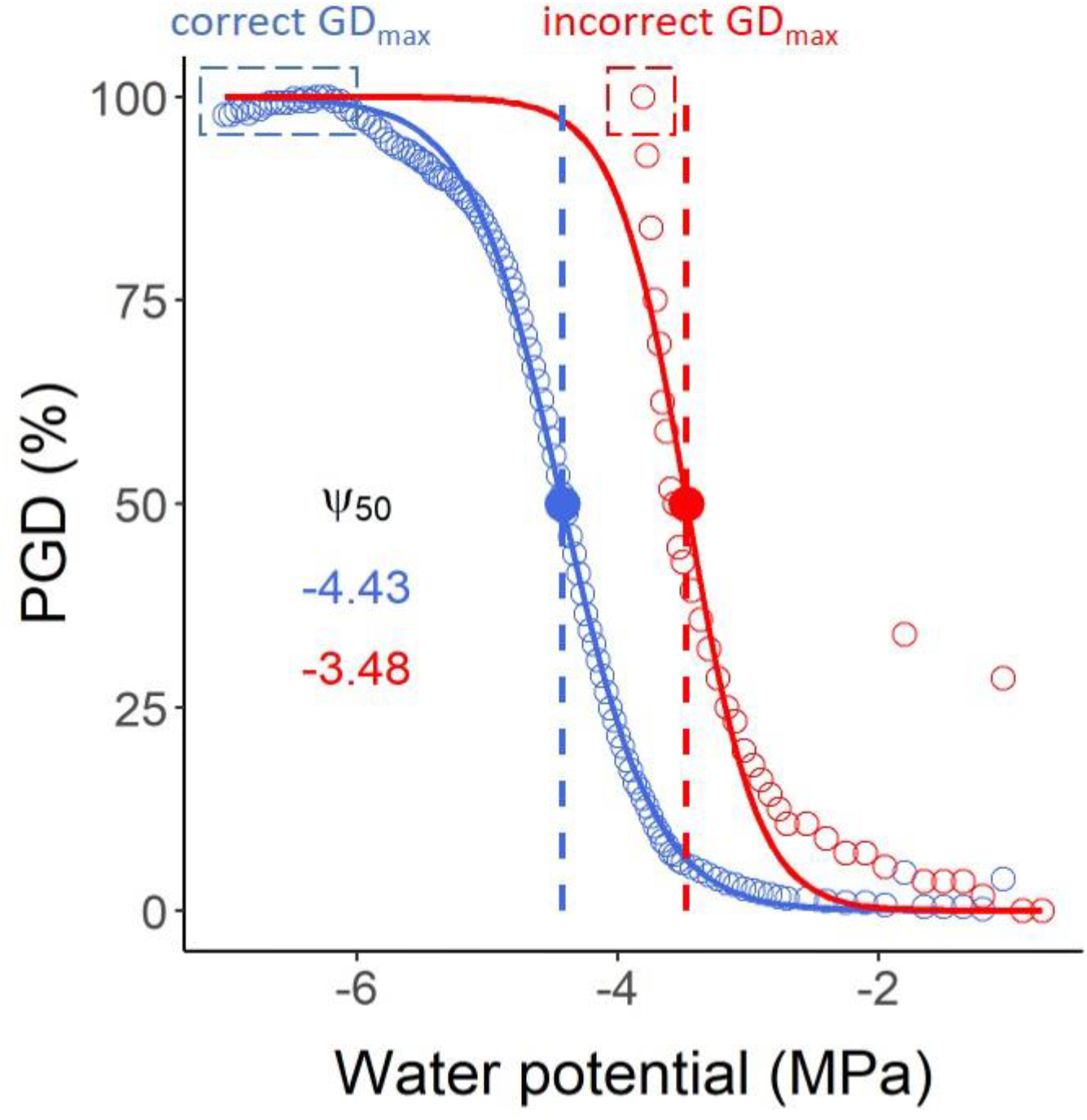
Vulnerability curves estimated with a Pneumatron for *Olea europea*, considering stable maximum gas discharge (GD_max_) measurements (a plateau below-6 MPa, in blue), and using a incorrect value as GD_max_ (in red). The water potential at which 50% of gas discharge was estimated (Ψ_50_, in MPa) is shown with the same colors. The red curve and its Ψ_50_ value should not be interpreted as accurate measurements due to incorrect estimation of GD_max_.

#### 3.2.8 Taking water potential measurements

When plotting vulnerability curves, xylem water potentials are required for the x-axis. Xylem water potential can be measured in different ways, with a pressure chamber or psychrometer.

##### 3.2.8.1 Water potential with pressure chamber

The branches need to be bagged at least 30 min before the measurements. If the whole branch is put in a dark environment (black plastic bag), the xylem water potential of a branch should be in equilibrium with the leaf water potential. Then, leaf water potential is assumed to be the xylem water potential when using the pressure chamber.

Excise one to two leaves from the branch and measure the leaf water potential using a pressure chamber. After cutting the leaves from the branch, it is important to apply glue to seal the cuts, because even cut petioles may contain vessels that run directly into the stem xylem and could lead to artificial air entry. Then, the branches are left to dehydrate and new measurements are taken. Write down the water potential results and the exact time of the measurement, so that water potential data and pneumatic measurements can be matched later.

Although the Pneumatron generally takes measurements at 15 min time intervals, it is practically not feasible to take water potential measurements with a pressure chamber at such high frequency. Therefore, estimating the decline in xylem water potential over time by interpolation is recommended (Pereira et al., 2020a). Generally, water potential measurements should be taken more frequently during the start of the dehydration process, and longer time periods can be taken once stomata have closed and a linear decline in water potential has been obtained. Thus, usually five to ten measurements are enough to estimate the water potential decline over the entire dehydration period, depending also on the leaf availability.

##### 3.2.8.2 Water potential with stem/leaf psychrometer

Leaf or stem psychrometers can be used, also in combination with pressure chamber measurements. The great advantage of using psychrometers, is that once they have been installed, the water potential is automatically monitored at a certain time interval, which could be set to the same time interval as the Pneumatron measurements.

## 4 Results

### 4.1 Data extraction and analysis of the vulnerability curves using the R script

The R script (https://github.com/Pneumatron/analysis) can be used to analyse the data. For this, the raw data from the Pneumatron (log.csv file) and another file with water potential data are needed, saved as a comma delimited file (csv). Headers of three columns must be defined: “date”, “hour”, and “psy” (date and hour, in the format of dd/mm/yyyy and hh:mm, respectively, and water potential measurements (psy)). Then, it is needed to input the experimental conditions as indicated at the beginning of the R script (file name and address, time for the initial and final pressures, reservoir volume, and atmospheric pressure). After running the script in R, the water potential between two consecutive measurements will be estimated, assuming a linear decrease over time. For example, from six measured points during two days of branch dehydration (with three measurements at two hour intervals during early dehydration, and three measurements every five to eight hours), it will estimate the water potential every 15 min, which corresponds to the same default time interval of Pneumatron measurements.

The script will also save three figures: the vulnerability curve (Fig. 6a), the volume of gas discharged versus water potential (Fig. 6b), and the volume of gas discharged versus time (Fig. 6c). By comparing the vulnerability curve (Fig. 6a) with the gas discharge curves (Fig. 6b and c), it is possible to infer problems regarding leakages or water potential measurements. Also, a file with the summarized results will be saved (results.csv). The script estimates the Ψ_50_, Ψ_88_, and Ψ_12_ by fitting a sigmoidal curve (p50.pad, p12.pad, and p88.pad results, following Pammenter and Vander Willigen (1998), or from the nearest data point measured directly by the Pneumatron (p50_near, p12_near, and p88_near results).

**Figure 6:**
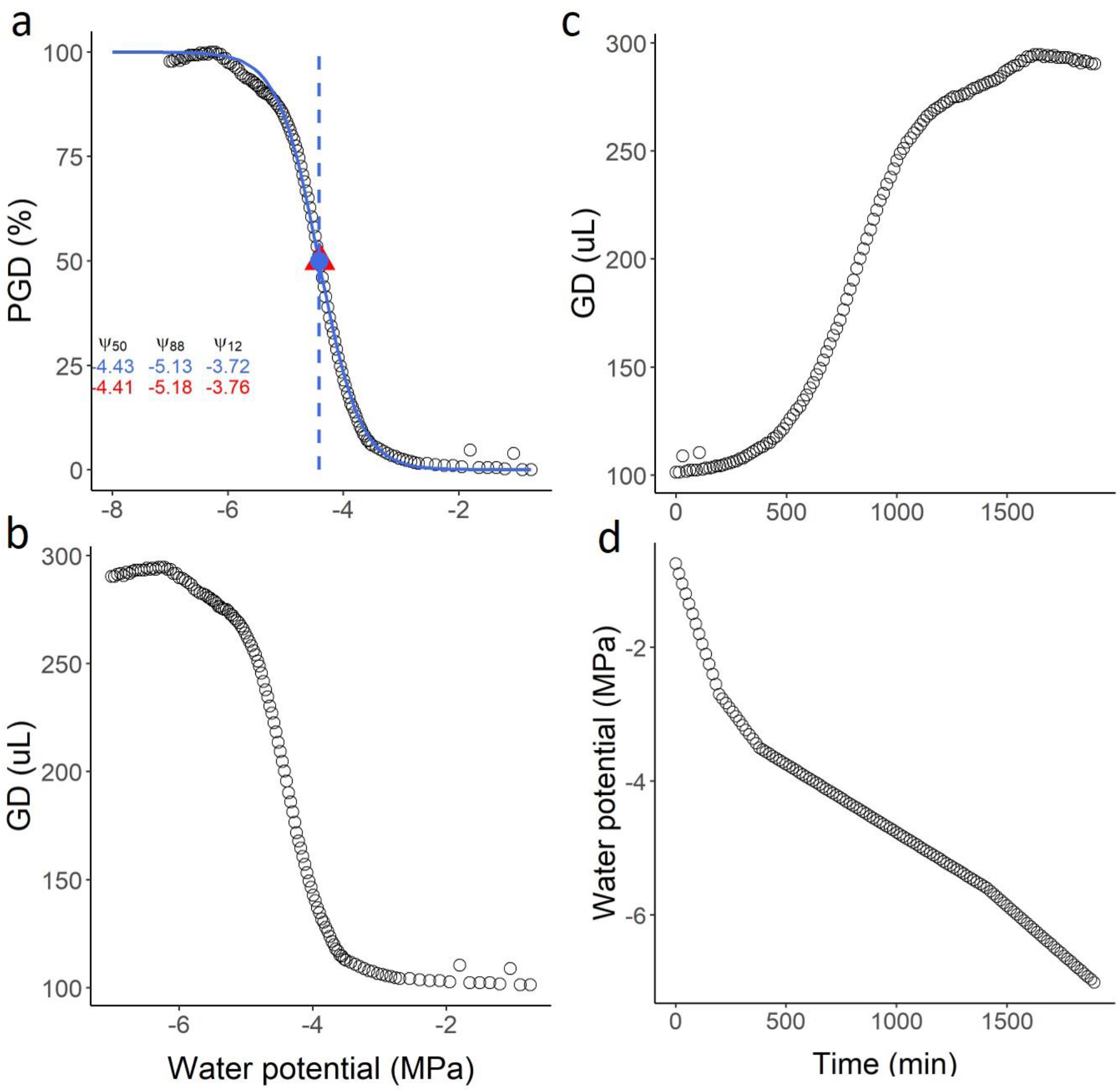
Example of output graphs from the R-script to analyse pneumatic gas discharge volumes (GD) measured with a Pneumatron for *Olea europea*. In (a) the vulnerability curve and the traits estimated from the sigmoidal curve (in blue) are shown. The values in red are the measured values at 12%, 50% and 88% PGD (percentage of gas discharge). The water potential corresponding to 50% PGD is shown in the graph as the estimated (blue circle) or measured value (red triangle). In (b) and (c) the gas discharged (raw data) is shown as a function of water potential (b) and time (c). In (d) the estimated water potential for every 15 min is shown as a function of time.

### 4.2 Vulnerability curves using an Excel spreadsheet

You may use the file data_example_pneumatron.xlsx to analyse the pneumatic data (Supplementary Material 2). A complete tutorial to use this excel file is shown in Supplementary Material 1. By curve fitting, vulnerability traits can be estimated in Excel, as shown in Supplementary Material 2.

### 4.3 Troubleshooting

During the construction and operation of Pneumatron devices, some of the following issues could be problematic. The most common problems are related to a leakage between the components, malfunctioning of data logger shield, or the solenoid valve. On the other hand, problems related to the Arduino board, electric components (such as Mosfets, resistors, etc.), or the vacuum pump are rare.

#### A. Nothing works

- Check the power supply.
- Check connections on the different shields and boards.
- Check that the pin connections are matching their counterparts.
- Check the pressure sensor and/or the ADC 16bits converter is accurately connected.
- Connect the Pneumatron to a computer, open the Arduino software and then open the serial monitor. Read the potential error messages if nothing appears to work (See section D).

#### B. Frequent or continuous pumping of the vacuum pump (leakage)

- If the main source of leakage has been identified to be the solenoid valve, consider replacing it.
- An alternative, but uncommon source of leakage can be the T junction.
- A leakage may also come from a not accurately inserted tube. Check if the solenoid valve and the pump are properly connected (following the correct orientation).
- The pump screws may be loose, and thus the pump cannot create the pressure declared in the software.

#### C. No pumping at all when powered

- Check if the pressure is near zero and not between 10 and 50 kPa. The solenoid valve may not open properly at the end of a measurement and the pressure does not return to the atmospheric pressure. Consider increasing the tubing volume between the pump and solenoid valve or replace the solenoid.
- Check the connection to the pump.
- Check if the pressure sensor and/or the ADC 16bits converter are correctly connected.
- Check if the programme has been properly uploaded on the board.

#### D. No data recorded or unreasonable values (all identical values or outside range values)

- SD card may be set on locked, corrupted, unformatted, or not fully inserted in the data logger shield. Change to the unlocked position, format the card, or check if the card is fully inserted and in the right orientation.
- Check the pressure sensor and/or the ADC 16bits converter is correctly connected.
- There could be a problem with the data logger shield. The pins may be not well connected, or another problem that is hard to detect may occur. Consider replacing it.
- An error in the recording of time can be due to a different RTC (Real Time Clock), The programme has been develop with the RTC DS1307, but check the data logger. If the inscription PCF8523 is shown, you need to change RTC_DS1307 to RTC_PCF8523, and change rtc.isrunning by rtc.initialized.

#### E. The pump seems to run under powered

- Check the power supply. The voltage or power might be too low. Consider replacing the power supply.

#### F. Arduino flashes rapidly but nothing else works

- This occurs when the voltage from the power supply is inferior to the optimal value. Consider changing the power supply.
- Alternatively, too many Mosfet transistors and/or too many pressure sensors are connected. Consider supplying the voltage to the Arduino from another source.

## 5 Discussion

The construction manual, the user manual details, and the software presented in this paper provide users with a Pneumatron device to estimate xylem embolism resistance. Vulnerability curves based on pneumatic measurements have been compared against various alternative methods (Pereira et al., 2016; Bittencourt et al., 2018; Zhang et al., 2018; Jansen et al., 2020). These earlier experiments showed that the pneumatic method provides accurate estimations of embolism resistance similar to the bench dehydration method, centrifuge-flow measurements (ChinaTron or cavitron; Cochard et al. (2005); Wang et al. (2014)), and the optical method (Brodribb et al., 2016). A comparison of Ψ_50_ data based on the Pneumatic method with alternative approaches for 50 angiosperm species shows a strong and highly significant correlation (R^2^ = 0.88, data not shown) considering published (Pereira et al., 2016, 2021; Zhang et al., 2018; Guan et al., 2020; Sergent et al., 2020) and unpublished results. Moreover, we used the Unit Pipe Pneumatic model to show that 91% of the gas volume extracted in 15 s of simulation comes from the first two series of intact and embolised vessels, while only 9% of the gas extracted results from xylem sap (Yang et al., submitted), which is saturated or supersaturated with gas (Schenk et al., 2016). This modelling exercise also showed that the Pneumatron may underestimate embolism resistance by 2 to 17%, with a typical measuring error of 0.11 MPa for Ψ_50_ values. Such measuring accuracy is better or at least equal to the typical measuring disagreement by various hydraulic methods based on the same species (Cochard et al., 2013; Jansen et al., 2015).

The high measuring accuracy of the Pneumatron is mainly due to the fast axial diffusion of gas across intervessel pit membranes (Kaack et al., 2019), while radial diffusion across cell walls and between the xylem and the bark is extremely slow (Sorz and Hietz, 2006; Wang et al., 2015). Even if intervessel pit membranes are hydrated, with a characteristic water volume fraction of ca. 80% (Zhang et al., 2020), diffusion across 200 to 1,000 nm thick pit membranes will be fast and within seconds (Yang et al., 2020). Considering the low costs, easy operation and fast analysis, the Pneumatron has considerable advantages over most conventional, both hydraulic and non-hydraulic methods (Table 1). The manual pneumatic approach, however, could result in a considerably larger measuring error due to unprecise recording of fast gas diffusion during the first seconds of gas extraction. Therefore, we recommend users to work with a Pneumatron device instead of the manual pneumatic approach, preferentially using the automated programme. In general, less data points are collected with the semi-automated mode and the connection and disconnection of samples may slightly change the leakage rates and tubing volume, which affects the measurement precision and the correct detection of the GD_min_ and GD_max_ plateaus (Fig. 4).

**Table 1:** Overview of the most common methods used to estimate xylem embolism resistance with the technical advantages, disadvantages, and key-references.

| Method | Advantage | Disadvantage | Key references |
| --- | --- | --- | --- |
| Bench dehydration | Cheap | Time-consuming; plenty of plant material needed; cutting-artefact; wounding artefact; destructive | (Sperry et al., 1988; Tyree et al., 1992; Bréda et al., 1993; Wheeler et al., 2013; De Baerdemaeker et al., 2019; Bonetti et al., 2020) |
| Centrifuge and flow-centrifuge | Fast; measurements rely on water transport under negative pressure | Open-vessel artefact; expensive; self-construction is technically highly challenging; hydraulic artefacts; destructive | (Alder et al., 1997; Cochard et al., 2005; Torres-Ruiz et al., 2014, 2017; Wang et al., 2014; Peng et al., 2019) |
| Air-injection double end | Relatively fast | Effervescence; vessel length exceeds cavitation chamber length; hydraulic artefacts; destructive | (Cochard et al., 1992; Gullo and Salleo, 1992; Yin and Cai, 2018) |
| Acoustic emissions | Rather expensive sensors; non-destructive | Data analysis time-consuming and not straightforward | (Milburn and Johnson, 1966; Tyree et al., 1984; Nolf et al., 2015) |
| Optical vulnerability | Cheap; applicable to leaf xylem and the outermost xylem of roots and stems; non-destructive | Time-consuming image analysis; mainly two-dimensional view of embolism | (Brodribb et al., 2016)<br>http://www.opensourceov.org/ |
| MicroCT | Non-destructive imaging | Extremely expensive; limited beam-time availability; time-consuming image analysis | (Choat et al., 2016; Klepsch et al., 2018) |
| Magnetic resonance imaging | Non-destructive imaging, quantification of flow | Low resolution; expensive | (Hochberg et al., 2016; Bouda et al., 2019) |
| Pneumatron | Cheap and straightforward operation, fast data analysis; 1 sample = 1 vulnerability curve | Adjustment of the discharge tube volume; may not be applicable to gymnosperms; destructive; no visual, anatomical information | (Pereira et al., 2016, 2020a; Bittencourt et al., 2018; Zhang et al., 2018) |

Yet, new users should be made well aware about the pneumatic principles, which are different from hydraulic measurements, and should be familiar with the basics that determine gas diffusion kinetics. These physical laws include the ideal gas law, Henry’s law for gas concentration partitioning between liquid and gas phases at equilibrium, and Fick’s law for diffusion. When the Pneumatic method is incorrectly applied, data are likely misinterpreted (Chen et al., 2020; Sergent et al., 2020; Pereira et al., 2021). The two most common mistakes include incorrect adjustment of the discharge tube volume, and the lack of stable gas discharge measurements at the beginning and the end of pneumatic measurements (GD_min_ and GD_max_). Since pneumatic vulnerability curves are normalised against GD_min_ and GD_max_, it is crucial to have reliable values for these two reference points (see Fig. 4). The consequences of not adjusting or incorrectly adjusting the volume of the discharge tube include a relatively high measuring error. When the discharge tube is too large or too small, the difference between the minimum and maximum air discharge volumes will be too small, resulting in relatively high measurement uncertainty (Jansen et al., 2020).

The main advantages of the Pneumatron device for constructing vulnerability curves are its low construction costs (< €100), the easy, automated operation, the fast analysis of pneumatic data, and its measuring accuracy. These issues are especially relevant when measurements on a large number of samples or species are desired. The device can also be used under remote field conditions, although water potential measurements may then become more challenging than pneumatic measurements. Vessel dimensions do not affect the precision of embolism resistance based on pneumatic measurements, as long as the discharge tube volume has been adjusted properly. This means that the Pneumatron can be applied to liana’s and species with long vessels. In fact, gas diffusion becomes more precise for wide vessels based on a Unit Pipe Pneumatic model (Yang et al., 2020). Ring-porous species, however, may need to be tested carefully because earlywood vessels in growth rings that are more than one year old are not only dysfunctional after one year, but also long and wide, and typically plugged with tyloses (Sano et al., 2011). One solution to work with ring-porous species is to glue off the xylem of previous growth rings (Zhang et al., 2018). Species that secrete resin, mucilage, oil or other substances, could also be problematic and should thus be tested carefully. Removing a thin slice at the cut-open xylem between measurements may be a solution.

A disadvantage is that the method is destructive and requires cutting open xylem tissue. Unlike hydraulic measurements, however, air entry of cut-open conduits should be aimed for as cut-open conduits function as an extension of the discharge tube. In case xylem samples that are collected would be under positive xylem pressure (Schenk et al., 2020), it is possible that xylem sap will come out at the cut-open end, which will not only be problematic for pneumatic measurements, but also for hydraulic methods. Other forms of wounding are likely slow and unlikely to affect the vessel dimensions during dehydration. Special attention should also be paid to non-xylem tissue such as pith, which may undergo some cracks and shrinkage during dehydration. Our experience, however, is that cracks or shrinkage are frequently not problematic when samples with relatively small pith proportions are used. Even if air from the pith tissue is extracted, this becomes only problematic when the gas volume extracted from the pith increases considerably, which is unlikely due to slow diffusion to this tissue. Also, cracks will only affect gas extraction amounts if these are connected to cut xylem conduits or the cut xylem area. Another disadvantage is that the method has not been successfully applied yet to gymnosperms. While the narrow tracheid dimensions could partially explain this, it is assumed that the torus-margo pit membranes of conifers could result in fast aspiration of the pit membrane (Zelinka et al., 2015; Schulte and Hacke, 2020), which prevents gas extraction from intact, embolised tracheids. It is possible, however, that a modification of the applied vacuum, discharge tube volume, and/or extraction time is needed to successfully apply the Pneumatron to gymnosperm xylem.

In addition to studies on xylem embolism, the Pneumatron has also been used to estimate vessel length distribution in an easy and much faster way than conventional methods (Williamson and Milburn, 2017; Link et al., 2018; Pereira et al., 2020b). This is possible by an easy modification of the reservoir volume and using the same software described here for the semi-automated mode. The data analysis is also straightforward by using a specific R-script, and thus, it may pave the way for fast measurements of vessel length distributions, and the non-random distribution of vessel ends near nodes, side branches, stem-petioles transitions, etc. Pneumatic estimations of the hydraulically weighted vessel length have been validated against the silicon-injection method for five species, and the air-injection method of Cohen et al., (2003) for seven species (Yang et al., submitted).

We hope this manual makes the Pneumatron accessible to many plant biologists as a high quality, low-cost, versatile tool. If required, the current version of the device can be modified and improved by using updated or alternative hardware and software. Alternatively, a commercial and user-friendly version is already available (Plantem – Plant and Environment Technologies, Campinas, Brazil). Different versions have been used satisfactorily so far, using other components than proposed here (Pereira et al., 2020a; Wu et al., 2020), or increasing the number of samples that are simultaneously measured (Pereira et al., 2020a). Any modification of the hardware, such as the changes needed to allow simultaneous measurements on multiple samples, will require further adaptation to the software as well as the R-script for data analysis. We encourage future users to share modified and updated versions with other users in a collaborative spirit as this would further improve pneumatic measurements of xylem and avoid misinterpretation of data.

## 6 Conflict of Interest

The authors declare that the research was conducted in the absence of any commercial or financial relationships that could be construed as a potential conflict of interest.

## 7 Author Contributions

CLT, LP, MTM, and XG wrote the first version of the manuscript. LP developed the R-script and the Arduino software, this latter with contributions from CLT and PRLB. CLT created the calibration protocol, calibration software, and the troubleshooting section. XG and RVR wrote the protocol and the template for the Excel analysis. The manuscript received substantial contributions from SJ, RVR, PRLB, and RO.

## 8 Funding

The authors acknowledge funding from the German Research Foundation (SJ, project nr. 383393940 and nr. 410768178) and from the São Paulo Research Foundation (RVR, MTM, and LP, Grant #2019/15276-8).

## 9 Acknowledgments

Financial support to SJ and XG is provided by a research grant from the German Research Foundation (project nr. 383393940 and nr. 410768178). The authors acknowledge the São Paulo Research Foundation (FAPESP, Brazil) for the research grant (RVR, LP and MTM, Grant 2019/15276-8), fellowship (LP and RVR, Grant 2017/14075-3) and scholarship (MTM and RVR, Grant 2018/09834-5). RVR are fellows of the National Council for Scientific and Technological Development (CNPq, Brazil).

## 11 Supplementary Material

**Supplementary Material 1** – Figure S1 – Schematic connection of the electronic components in a Pneumatron device; Table S1 – Component list; Tutorial to use the excel file for data analysis.

**Supplementary Material 2** – Excel file for data analysis.

## 12 Data Availability Statement

The datasets generated for this study can be found in the Github https://github.com/Pneumatron.

## Notes

### Competing Interest Statement

The authors have declared no competing interest.

## References

Alder, N. N., Pockman, W. T., Sperry, J. S., and Nuismer, S. (1997). Use of centrifugal force in the study of xylem cavitation. J. Exp. Bot. 48, 665–674. doi:10.1093/jxb/48.3.665.

Bittencourt, P., Pereira, L., and Oliveira, R. (2018). Pneumatic method to measure plant xylem embolism. Bio-Protocol 8, 1–14. doi:10.21769/bioprotoc.3059.

Bonetti, S., Breitenstein, D., Fatichi, S., Domec, J. C., and Or, D. (2020). Persistent decay of fresh xylem hydraulic conductivity varies with pressure gradient and marks plant responses to injury. Plant Cell Environ., 1–16. doi:10.1111/pce.13893.

Bouda, M., Windt, C. W., McElrone, A. J., and Brodersen, C. R. (2019). In vivo pressure gradient heterogeneity increases flow contribution of small diameter vessels in grapevine. Nat. Commun. 10, 5645. doi:10.1038/s41467-019-13673-6.

Bréda, N., Cochard, H., Dreyer, E., and Granier, A. (1993). Field comparison of transpiration, stomatal conductance and vulnerability to cavitation of Quercus petraea and Quercus robur under water stress. Ann. des Sci. For. 50, 571–582. doi:10.1051/forest:19930606.

Brodribb, T. J., Skelton, R. P., Mcadam, S. A. M., Bienaimé, D., Lucani, C. J., and Marmottant, P. (2016). Visual quantification of embolism reveals leaf vulnerability to hydraulic failure. New Phytol. 209, 1403–1409. doi:10.1111/nph.13846.

Chen, Y.-J., Maenpuen, P., Zhang, Y.-J., Barai, K., Katabuchi, M., Gao, H., et al. (2020). Quantifing vulnerability to embolism in tropical trees and lianas using five methods: Can discrepancies be explained by xylem structural traits?. doi:10.1111/nph.16927.

Choat, B., Badel, E., Burlett, R., Delzon, S., Cochard, H., and Jansen, S. (2016). Noninvasive measurement of vulnerability to drought-induced embolism by X-Ray microtomography. Plant Physiol. 170, 273–282. doi:10.1104/pp.15.00732.

Cochard, H., Badel, E., Herbette, S., Delzon, S., Choat, B., and Jansen, S. (2013). Methods for measuring plant vulnerability to cavitation: A critical review. J. Exp. Bot. 64, 4779–4791. doi:10.1093/jxb/ert193.

Cochard, H., Cruiziat, P., and Tyree, M. T. (1992). Use of positive pressures to establish vulnerability curves: Further support for the air-seeding hypothesis and implications for pressure-volume analysis. Plant Physiol. 100, 205–209. doi:10.1104/pp.100.1.205.

Cochard, H., Damour, G., Bodet, C., Tharwat, I., Poirier, M., and Améglio, T. (2005). Evaluation of a new centrifuge technique for rapid generation of xylem vulnerability curves. Physiol. Plant. 124, 410–418. doi:10.1111/j.1399-3054.2005.00526.x.

Cohen, S., Bennink, J., and Tyree, M. (2003). Air method measurements of apple vessel length distributions with improved apparatus and theory. J. Exp. Bot. 54, 1889–1897. doi:10.1093/jxb/erg202.

De Baerdemaeker, N. J. F., Arachchige, K. N. R., Zinkernagel, J., Van Den Bulcke, J., Van Acker, J., Schenk, H. J., et al. (2019). The stability enigma of hydraulic vulnerability curves: Addressing the link between hydraulic conductivity and drought-induced embolism. Tree Physiol. 39, 1646–1664. doi:10.1093/treephys/tpz078.

Greenidge, K. N. H. (1952). An approach to the study of vessel length in hardwood species. Am. J. Bot. 39, 570–574. doi:10.1002/j.1537-2197.1952.tb13070.x.

Guan, X., Pereira, L., McAdam, S., Cao, K.-F., and Jansen, S. (2020). No gas source, no problem: pre-existing embolism may affect non-pressure driven embolism spreading in angiosperm xylem by gas diffusion. Authorea Prepr., 1–18. doi:10.22541/au.159674582.24283294.

Gullo, M. A., and Salleo, S. (1992). Water storage in the wood and xylem cavitation in 1-year-old twigs of Populus deltoides Bartr. Plant, Cell Environ. 15, 431–438. doi:10.1111/j.1365-3040.1992.tb00993.x.

Hochberg, U., Albuquerque, C., Rachmilevitch, S., Cochard, H., David-Schwartz, R., Brodersen, C. R., et al. (2016). Grapevine petioles are more sensitive to drought induced embolism than stems: evidence from in vivo MRI and microcomputed tomography observations of hydraulic vulnerability segmentation. Plant. Cell Environ. 39, 1886–1894. doi:10.1111/pce.12688.

Jansen, S., Guan, X., Kaack, L., Trabi, C.L., Miranda, M. T., Ribeiro, R. V, et al. (2020). The Pneumatron estimates xylem embolism resistance in angiosperms based on gas diffusion kinetics: a mini-review. Acta Hortic. in press.

Jansen, S., Schuldt, B., and Choat, B. (2015). Current controversies and challenges in applying plant hydraulic techniques. New Phytol. 205, 961–964. doi:10.1111/nph.13229.

Kaack, L., Altaner, C. M., Carmesin, C., Diaz, A., Holler, M., Kranz, C., et al. (2019). Function and three-dimensional structure of intervessel pit membranes in angiosperms: a review. IAWA J. 40, 673–702. doi:10.1163/22941932-40190259.

Klepsch, M., Zhang, Y., Kotowska, M. M., Lamarque, L. J., Nolf, M., Schuldt, B., et al. (2018). Is xylem of angiosperm leaves less resistant to embolism than branches? Insights from microCT, hydraulics, and anatomy. J. Exp. Bot. 69, 5611–5623. doi:10.1093/jxb/ery321.

Link, R. M., Schuldt, B., Choat, B., Jansen, S., and Cobb, A. R. (2018). Maximum-likelihood estimation of xylem vessel length distributions. J. Theor. Biol. 455, 329–341. doi:10.1016/j.jtbi.2018.07.036.

Milburn, J., and Johnson, R. (1966). The conduction of sap. II. Detection of vibrations produced by sap cavitation in Ricinus xylem. Planta 69, 43–52.

Nolf, M., Beikircher, B., Rosner, S., Nolf, A., and Mayr, S. (2015). Xylem cavitation resistance can be estimated based on time-dependent rate of acoustic emissions. New Phytol. 208, 625–632. doi:10.1111/nph.13476.

Pammenter, N. W., and Van der Willigen, C. (1998). A mathematical and statistical analysis of the curves illustrating vulnerability of xylem to cavitation. Tree Physiol. 18, 589–593. doi:10.1093/treephys/18.8-9.589.

Peng, G., Yang, D., Liang, Z., Li, J., and Tyree, M. T. (2019). An improved centrifuge method for determining water extraction curves and vulnerability curves in the long-vessel species Robinia pseudoacacia. J. Exp. Bot. 70, 4865–4875. doi:10.1093/jxb/erz206.

Pereira, L., Bittencourt, P. R. L., Oliveira, R. S., Junior, M. B. M., Barros, F. V, Ribeiro, R. V., et al. (2016). Plant pneumatics: stem air flow is related to embolism – new perspectives on methods in plant hydraulics. New Phytol. 211, 357–370. doi:10.1111/nph.13905.

Pereira, L., Bittencourt, P. R. L., Pacheco, V. S., Miranda, M. T., Zhang, Y., Oliveira, R. S., et al. (2020a). The Pneumatron: An automated pneumatic apparatus for estimating xylem vulnerability to embolism at high temporal resolution. Plant. Cell Environ. 43, 131–142. doi:10.1111/pce.13647.

Pereira, L., Bittencourt, P. R. L., Rowland, L., Brum, M., Miranda, M. T., Pacheco, V. S., et al. (2021). Using the Pneumatic method to estimate embolism resistance in species with long vessels: A commentary on the article “A comparison of five methods to assess embolism resistance in trees.”For. Ecol. Manage. 479, 118547. doi:10.1016/j.foreco.2020.118547.

Pereira, L., Miranda, M. T., Pires, G. S., Pacheco, V. S., Guan, X., Kaack, L., et al. (2020b). A semi-automated method for measuring xylem vessel length distribution. Theor. Exp. Plant Physiol. doi:10.1007/s40626-020-00189-4.

Sano, Y., Morris, H., Shimada, H., Ronse De Craene, L. P., and Jansen, S. (2011). Anatomical features associated with water transport in imperforate tracheary elements of vessel-bearing angiosperms. Ann. Bot. 107, 953–964. doi:10.1093/aob/mcr042.

Schenk, H. J., Espino, S., Visser, A., and Esser, B. K. (2016). Dissolved atmospheric gas in xylem sap measured with membrane inlet mass spectrometry. Plant Cell Environ. 39, 944–950. doi:10.1111/pce.12678.

Schenk, H. J., Jansen, S., and Hölttä, T. (2020). Positive pressure in xylem and its role in hydraulic function. New Phytol., nph.17085. doi:10.1111/nph.17085.

Schulte, P. J., and Hacke, U. G. (2020). Solid mechanics of the torus-margo in conifer inter-tracheid bordered pits. New Phytol. doi:10.1111/nph.16949.

Sergent, A. S., Varela, S. A., Barigah, T. S., Badel, E., Cochard, H., Dalla-Salda, G., et al. (2020). A comparison of five methods to assess embolism resistance in trees. For. Ecol. Manage. 468, 118175. doi:10.1016/j.foreco.2020.118175.

Sorz, J., and Hietz, P. (2006). Gas diffusion through wood: Implications for oxygen supply. Trees - Struct. Funct. 20, 34–41. doi:10.1007/s00468-005-0010-x.

Sperry, J. S., Donnelly, J. R., and Tyree, M. T. (1988). A method for measuring hydraulic conductivity and embolism in xylem. Plant, Cell Environ. 11, 35–40. doi:10.1111/j.1365-3040.1988.tb01774.x.

Torres-Ruiz, J. M., Cochard, H., Choat, B., Jansen, S., López, R., Tomášková, I., et al. (2017). Xylem resistance to embolism: presenting a simple diagnostic test for the open vessel artefact. New Phytol. 215, 489–499. doi:10.1111/nph.14589.

Torres-Ruiz, J. M., Cochard, H., Mayr, S., Beikircher, B., Diaz-Espejo, A., Rodriguez-Dominguez, C. M., et al. (2014). Vulnerability to cavitation in Olea europaea current-year shoots: further evidence of an open-vessel artifact associated with centrifuge and air-injection techniques. Physiol. Plant. 152, 465–474. doi:10.1111/ppl.12185.

Tyree, M. T., Alexander, J., and Machado, J.-L. (1992). Loss of hydraulic conductivity due to water stress in intact juveniles of Quercus rubra and Populus deltoides. Tree Physiol. 10, 411–415. doi:10.1093/treephys/10.4.411.

Tyree, M. T., Dixon, M. A., Tyree, E. L., and Johnson, R. (1984). Ultrasonic Acoustic Emissions from the Sapwood of Cedar and Hemlock. Plant Physiol. 75, 988–992. doi:10.1104/pp.75.4.988.

Wang, Y., Burlett, R., Feng, F., and Tyree, M. T. (2014). Improved precision of hydraulic conductance measurements using a Cochard rotor in two different centrifuges. J. Plant Hydraul. 1, e007. doi:10.20870/jph.2014.e007.

Wang, Y., Liu, J., and Tyree, M. T. (2015). Stem hydraulic conductivity depends on the pressure at which it is measured and how this dependence can be used to assess the tempo of bubble pressurization in recently cavitated vessels. Plant Physiol. 169, 2597–2607. doi:10.1104/pp.15.00875.

Wheeler, J. K., Huggett, B. A., Tofte, A. N., Rockwell, F. E., and Holbrook, N. M. (2013). Cutting xylem under tension or supersaturated with gas can generate PLC and the appearance of rapid recovery from embolism. Plant, Cell Environ. 36, 1938–1949. doi:10.1111/pce.12139.

Williamson, V. G., and Milburn, J. A. (2017). Xylem vessel length and distribution: Does analysis method matter? A study using Acacia. Aust. J. Bot. 65, 292–303. doi:10.1071/BT16220.

Wu, M., Zhang, Y., Oya, T., Marcati, C. R., Pereira, L., and Jansen, S. (2020). Root xylem in three woody angiosperm species is not more vulnerable to embolism than stem xylem. Plant Soil 450, 479–495. doi:10.1007/s11104-020-04525-0.

Yang, D., Pereira, L., Peng1, G., Ribeiro, R. V., Kaack, L., Jansen, S., et al. (2020). A Unit Pipe Pneumatic model to simulate gas kinetics during measurements of embolism in excised angiosperm xylem. submitted.

Yin, P., and Cai, J. (2018). New possible mechanisms of embolism formation when measuring vulnerability curves by air injection in a pressure sleeve. Plant. Cell Environ. 41, 1361–1368. doi:10.1111/pce.13163.

Zelinka, S. L., Bourne, K. J., Hermanson, J. C., Glass, S. V., Costa, A., and Wiedenhoeft, A. C. (2015). Force-displacement measurements of earlywood bordered pits using a mesomechanical tester. Plant, Cell Environ. 38, 2088–2097. doi:10.1111/pce.12532.

Zhang, Y., Carmesin, C., Kaack, L., Klepsch, M. M., Kotowska, M., Matei, T., et al. (2020). High porosity with tiny pore constrictions and unbending pathways characterize the 3D structure of intervessel pit membranes in angiosperm xylem. Plant Cell Environ. 43, 116–130. doi:10.1111/pce.13654.

Zhang, Y., Lamarque, L. J., Torres-Ruiz, J. M., Schuldt, B., Karimi, Z., Li, S., et al. (2018). Testing the plant pneumatic method to estimate xylem embolism resistance in stems of temperate trees. Tree Physiol. 38, 1016–1025. doi:10.1093/treephys/tpy015.

