## Supplementary Material 1 for "A user manual to measure gas diffusion kinetics in plants: Pneumatron construction, operation and data analysis"

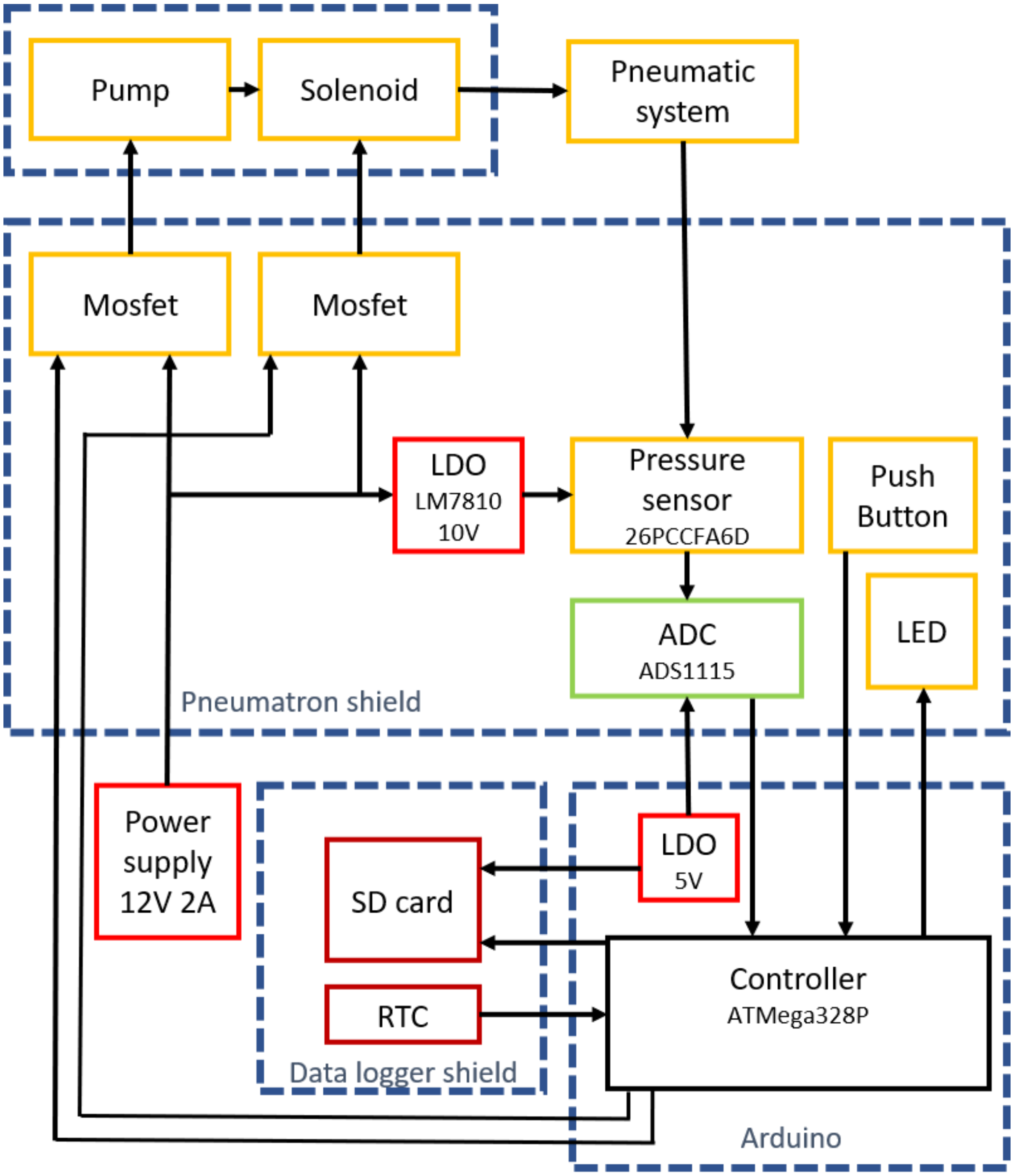

**Figure S1 –** Schematic connection of the electronic components in a Pneumatron device.

**Table S1** – Component list

| | Component | Manufacturer | Catalog/part code | Key specifications |
|---|---|---|---|---|
| 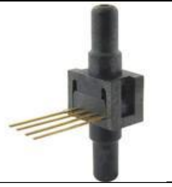 | Differential pressure sensor | Honeywell | 26PCCFA6D | 15 Psi<br><br>100 mV spam |
| 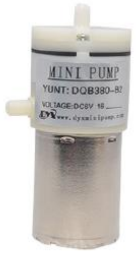 | Mini vacuum pump | Dyx, Shenzhen, China | DQB380-FB2 | 12 V, 700mA maximum |
| 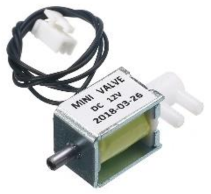 | Three-way solenoid valve<br><br>(12 volts) | Dongguan City-Electric Co., Dongguan, China | Fa0520F | 12 V, 500 mA maximum |
| 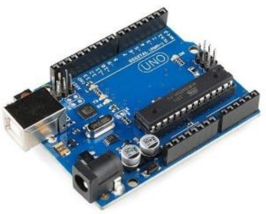 | Arduino UNO | | | |
| 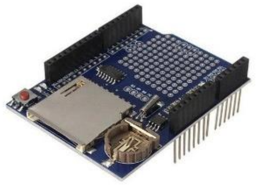 | Data logger shield | | | DS1307<br><br>MicroSD adapter plus level shifters |
| 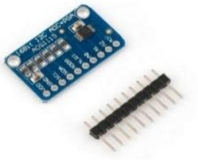 | Analogic to digital converter 16 bits | | ADS1115 breakout board | Configured to address 0x48 |

| | | | | |
|---|---|---|---|---|
| 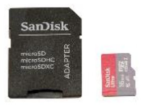 | Micro-SD memory card and SD adapter (see data logger module) | | | SD or SDHC |
| 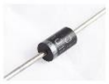 | 2 Schotksy Diode | | 1N4007 | Used as flyback diode; fast response, low forward voltage. |
| 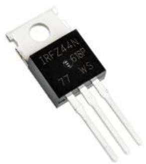 | 2 N-ChannelMOSFET | | IRFZ44NPBF | 5 V logic level; Igs >= 3A |
| 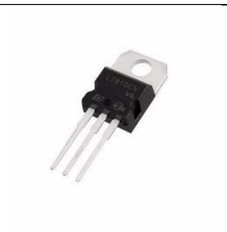 | Voltage regulator | | L7810 | |
| 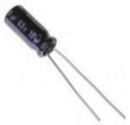 | Eletrolytic Capacitor 10UF 63V | | | |
| 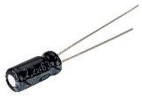 | Eletrolytic Capacitor 2.2UF 63V | | | |
| 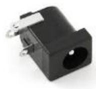 | DC Power Supply Jack Socket Female Panel Mount Connector | | | |
| 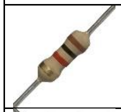 | Carbon resistor 1K | | | |
| 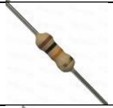 | Carbon resistor 10K | | | |
| 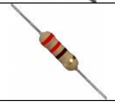 | Carbon resistor 220 ohms | | | |
| 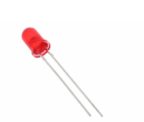 | Light Emitting Diode | | | |

| | | | | |
|---|---|---|---|---|
| 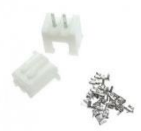 | Pin header, housing and cable crimp | | JST-XH | Vertical, 2.5 mm pitch (if XH) |
| 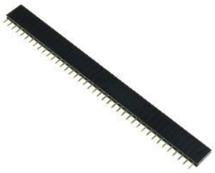 | Single Row Straight Female Pin Receptacle | | | 2.5 mm pitch |
| 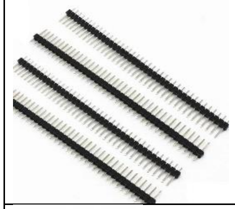 | Single Row Straight Male Pin Header Strip | | | |
| 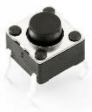 | Push button | | | 6 x 6 mm, normal open |
| 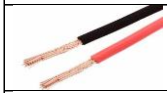 | Wires (red and black) | 20 cm each | | AWG <=28 |
| 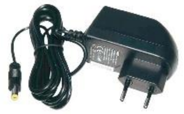 | Power source | | | 12 VDC, 2 A |
| 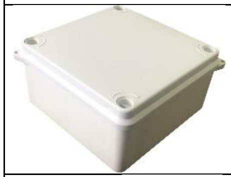 | Plastic enclosure | | | 10x10x5 cm |
| 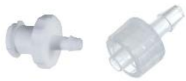 | Adapter Luers | Cole-Parmer | EW-30800-06 and EW-30800-24 | |
| 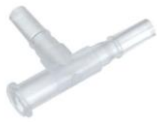 | Luer Adapter Tees, Male x Male x Female | Cole-Parmer | GY-45508-75 | |
| 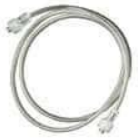 | Rigid Tubing | Cole-Parmer | EW-30600-62 | |
| 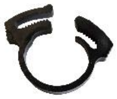 | Plastic clamps | Cole-Parmer | RZ-06832-02 | |

| | |
|---|---|
| 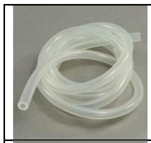 | Silicone tubing (2 mm Internal Diameter) |
| 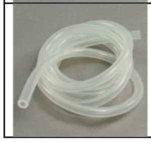 | Silicone tubing (4 mm ID and 6 mm OD) |

**Tutorial to use the excel file for data analysis.**

1.1 Get the data.

1.1 Import the csv file into Excel software.

1.2 Rearrange the data: Split the single column to separate ones.

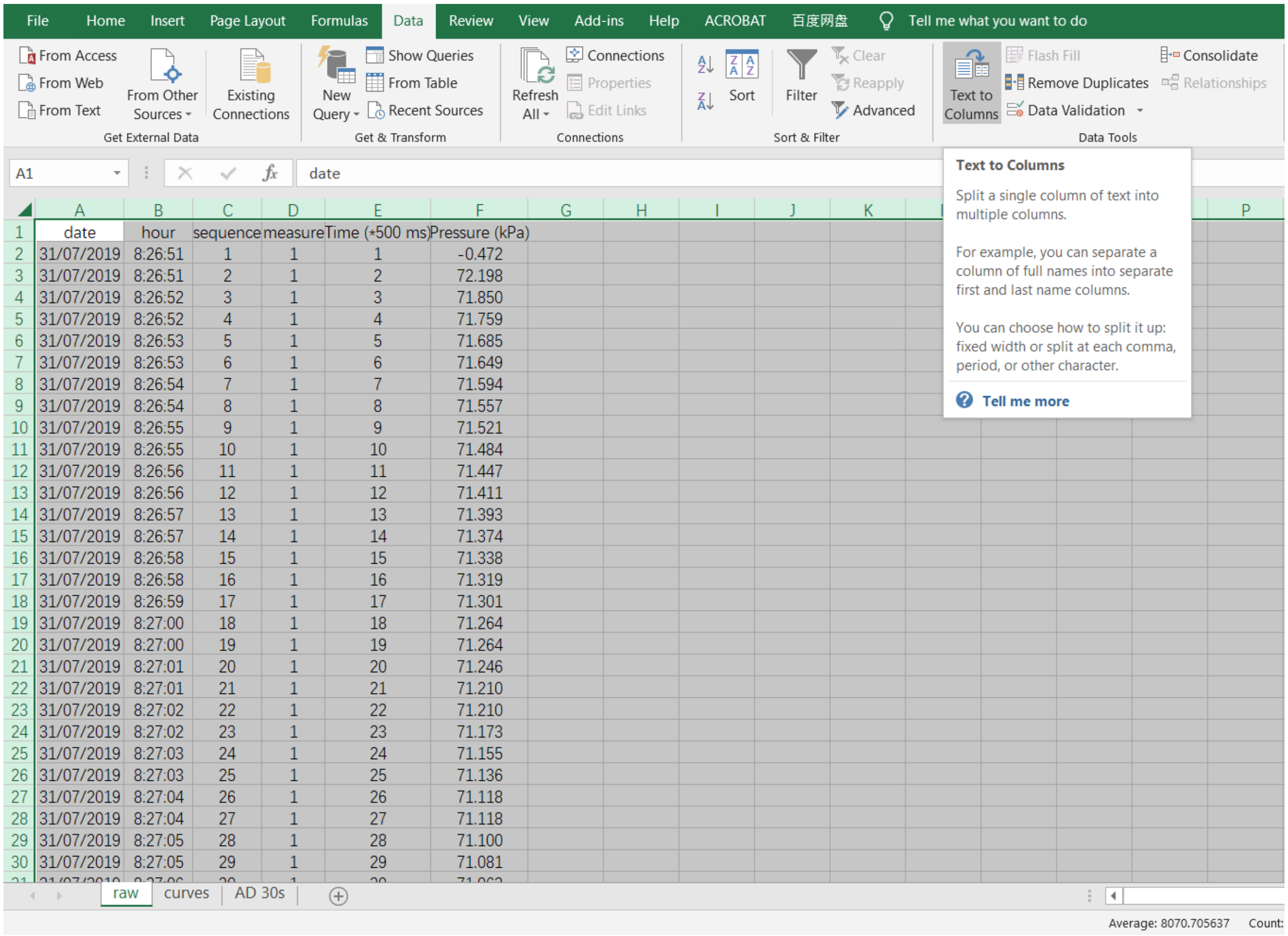

| | A | B | C | D | E | F |
|---|---|---|---|---|---|---|
| 1 | date | hour | sequence | measure | Time (*500 ms) | Pressure (kPa) |
| 2 | 31/07/2019 | 8:26:51 | 1 | 1 | 1 | -0.472 |
| 3 | 31/07/2019 | 8:26:51 | 2 | 1 | 2 | 72.198 |
| 4 | 31/07/2019 | 8:26:52 | 3 | 1 | 3 | 71.850 |
| 5 | 31/07/2019 | 8:26:52 | 4 | 1 | 4 | 71.759 |
| 6 | 31/07/2019 | 8:26:53 | 5 | 1 | 5 | 71.685 |
| 7 | 31/07/2019 | 8:26:53 | 6 | 1 | 6 | 71.649 |
| 8 | 31/07/2019 | 8:26:54 | 7 | 1 | 7 | 71.594 |
| 9 | 31/07/2019 | 8:26:54 | 8 | 1 | 8 | 71.557 |
| 10 | 31/07/2019 | 8:26:55 | 9 | 1 | 9 | 71.521 |
| 11 | 31/07/2019 | 8:26:55 | 10 | 1 | 10 | 71.484 |
| 12 | 31/07/2019 | 8:26:56 | 11 | 1 | 11 | 71.447 |
| 13 | 31/07/2019 | 8:26:56 | 12 | 1 | 12 | 71.411 |
| 14 | 31/07/2019 | 8:26:57 | 13 | 1 | 13 | 71.393 |
| 15 | 31/07/2019 | 8:26:57 | 14 | 1 | 14 | 71.374 |
| 16 | 31/07/2019 | 8:26:58 | 15 | 1 | 15 | 71.338 |
| 17 | 31/07/2019 | 8:26:58 | 16 | 1 | 16 | 71.319 |
| 18 | 31/07/2019 | 8:26:59 | 17 | 1 | 17 | 71.301 |
| 19 | 31/07/2019 | 8:27:00 | 18 | 1 | 18 | 71.264 |
| 20 | 31/07/2019 | 8:27:00 | 19 | 1 | 19 | 71.264 |
| 21 | 31/07/2019 | 8:27:01 | 20 | 1 | 20 | 71.246 |
| 22 | 31/07/2019 | 8:27:01 | 21 | 1 | 21 | 71.210 |
| 23 | 31/07/2019 | 8:27:02 | 22 | 1 | 22 | 71.210 |
| 24 | 31/07/2019 | 8:27:02 | 23 | 1 | 23 | 71.173 |
| 25 | 31/07/2019 | 8:27:03 | 24 | 1 | 24 | 71.155 |
| 26 | 31/07/2019 | 8:27:03 | 25 | 1 | 25 | 71.136 |
| 27 | 31/07/2019 | 8:27:04 | 26 | 1 | 26 | 71.118 |
| 28 | 31/07/2019 | 8:27:04 | 27 | 1 | 27 | 71.118 |
| 29 | 31/07/2019 | 8:27:05 | 28 | 1 | 28 | 71.100 |
| 30 | 31/07/2019 | 8:27:05 | 29 | 1 | 29 | 71.081 |
| [illegible] | [illegible] | [illegible] | [illegible] | [illegible] | [illegible] | [illegible] |

**Supplementary Figure S2:** Rearranging data

1.3 Add titles to column D, E and F with “Measure”, “Time (*500 ms)” and “Pressure (kPa)”

“Measure” gives the information of the exact round of measurement that is taking, “Time” tells the sequence of records during one measurement, and “Pressure” tells the corresponding pressure, which is recorded at every 500 ms.

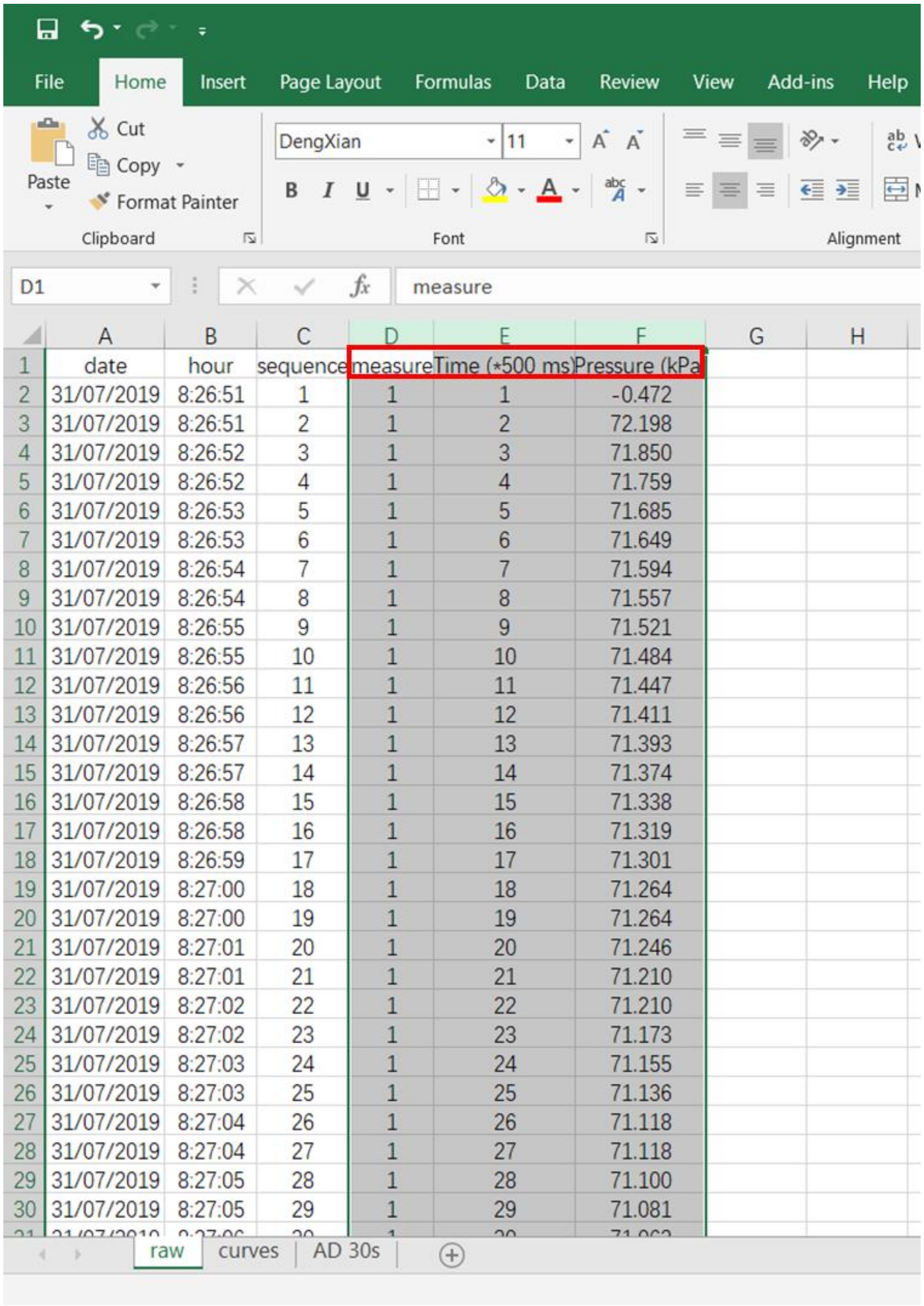

| | A | B | C | D | E | F |
|---|---|---|---|---|---|---|
| 1 | date | hour | sequence | measure | Time (*500 ms) | Pressure (kPa |
| 2 | 31/07/2019 | 8:26:51 | 1 | 1 | 1 | -0.472 |
| 3 | 31/07/2019 | 8:26:51 | 2 | 1 | 2 | 72.198 |
| 4 | 31/07/2019 | 8:26:52 | 3 | 1 | 3 | 71.850 |
| 5 | 31/07/2019 | 8:26:52 | 4 | 1 | 4 | 71.759 |
| 6 | 31/07/2019 | 8:26:53 | 5 | 1 | 5 | 71.685 |
| 7 | 31/07/2019 | 8:26:53 | 6 | 1 | 6 | 71.649 |
| 8 | 31/07/2019 | 8:26:54 | 7 | 1 | 7 | 71.594 |
| 9 | 31/07/2019 | 8:26:54 | 8 | 1 | 8 | 71.557 |
| 10 | 31/07/2019 | 8:26:55 | 9 | 1 | 9 | 71.521 |
| 11 | 31/07/2019 | 8:26:55 | 10 | 1 | 10 | 71.484 |
| 12 | 31/07/2019 | 8:26:56 | 11 | 1 | 11 | 71.447 |
| 13 | 31/07/2019 | 8:26:56 | 12 | 1 | 12 | 71.411 |
| 14 | 31/07/2019 | 8:26:57 | 13 | 1 | 13 | 71.393 |
| 15 | 31/07/2019 | 8:26:57 | 14 | 1 | 14 | 71.374 |
| 16 | 31/07/2019 | 8:26:58 | 15 | 1 | 15 | 71.338 |
| 17 | 31/07/2019 | 8:26:58 | 16 | 1 | 16 | 71.319 |
| 18 | 31/07/2019 | 8:26:59 | 17 | 1 | 17 | 71.301 |
| 19 | 31/07/2019 | 8:27:00 | 18 | 1 | 18 | 71.264 |
| 20 | 31/07/2019 | 8:27:00 | 19 | 1 | 19 | 71.264 |
| 21 | 31/07/2019 | 8:27:01 | 20 | 1 | 20 | 71.246 |
| 22 | 31/07/2019 | 8:27:01 | 21 | 1 | 21 | 71.210 |
| 23 | 31/07/2019 | 8:27:02 | 22 | 1 | 22 | 71.210 |
| 24 | 31/07/2019 | 8:27:02 | 23 | 1 | 23 | 71.173 |
| 25 | 31/07/2019 | 8:27:03 | 24 | 1 | 24 | 71.155 |
| 26 | 31/07/2019 | 8:27:03 | 25 | 1 | 25 | 71.136 |
| 27 | 31/07/2019 | 8:27:04 | 26 | 1 | 26 | 71.118 |
| 28 | 31/07/2019 | 8:27:04 | 27 | 1 | 27 | 71.118 |
| 29 | 31/07/2019 | 8:27:05 | 28 | 1 | 28 | 71.100 |
| 30 | 31/07/2019 | 8:27:05 | 29 | 1 | 29 | 71.081 |
| [illegible] | [illegible] | [illegible] | [illegible] | [illegible] | [illegible] | [illegible] |

**Supplementary Figure S3:** Adding headers

**1.4** Select the three columns named “Measure”, “Duration”, “Pressure”. Go to “Insert” and choose “PivotTable”.

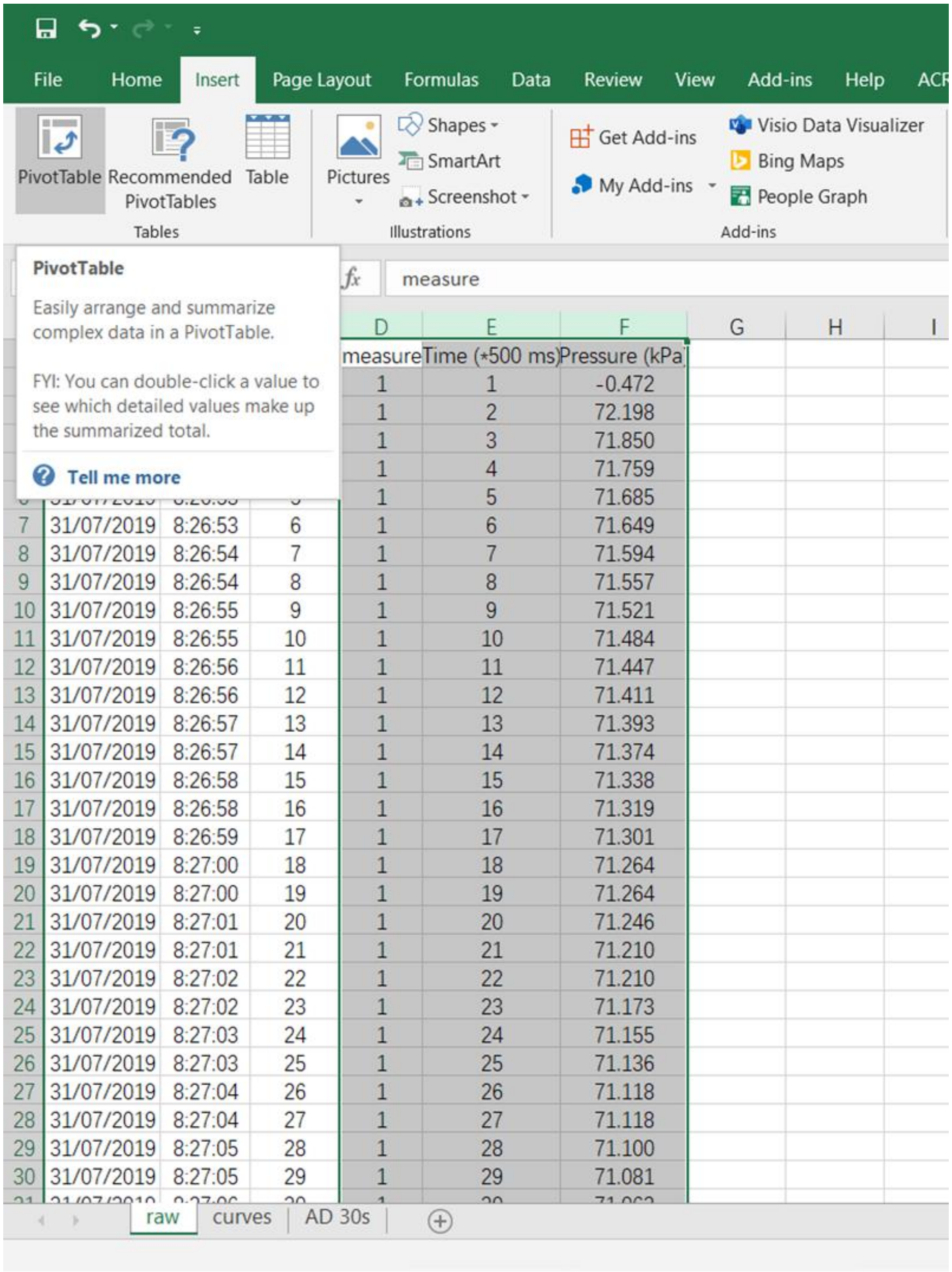

**Supplementary Figure S4:** Rearranging columns

1.5 Create a new sheet and drag columns.

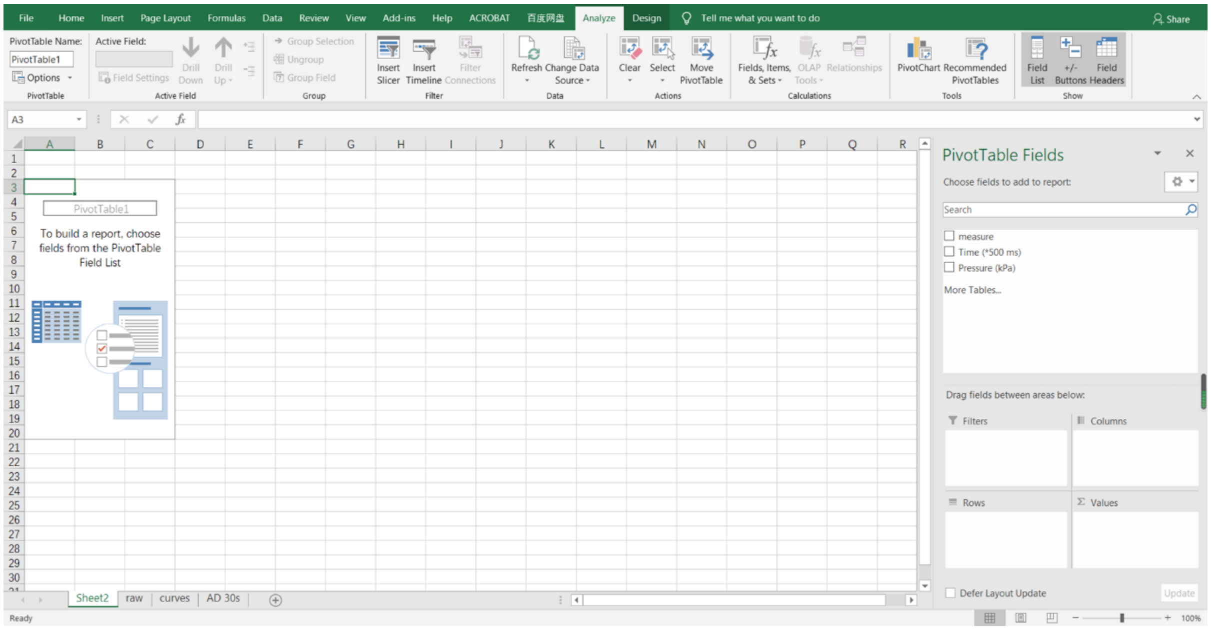

**Supplementary Figure S5:** New sheet

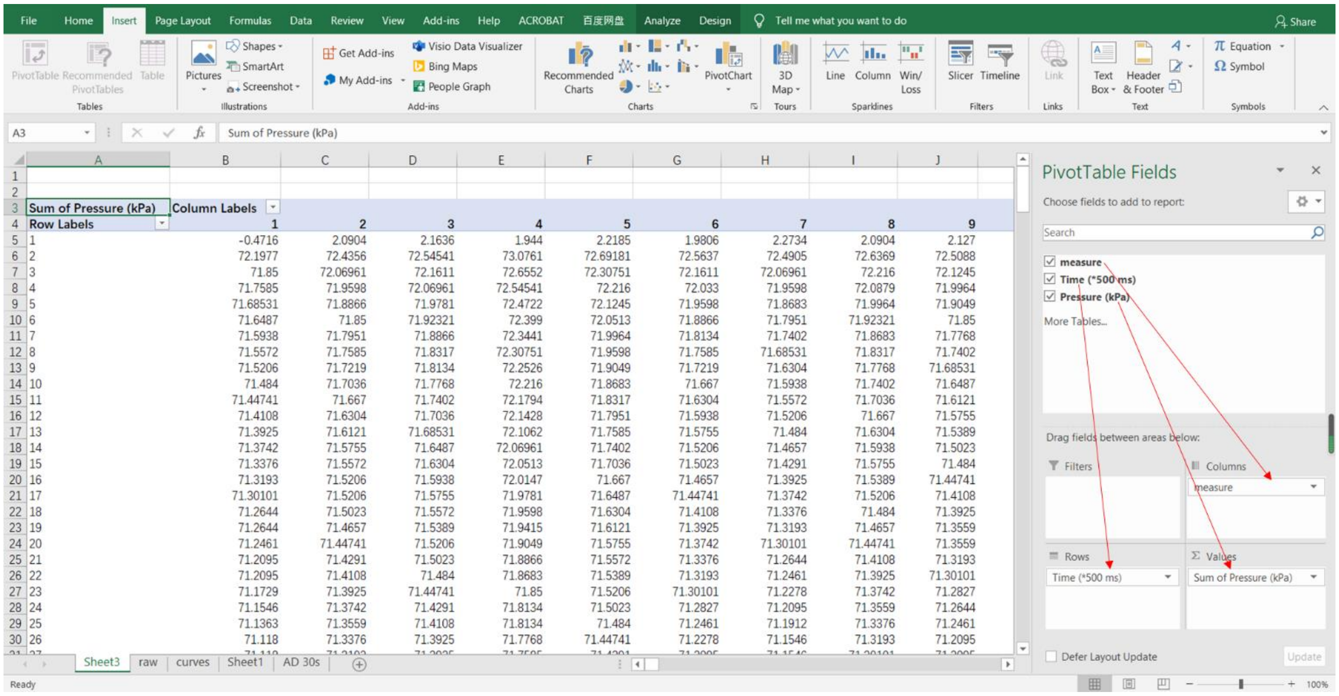

**Supplementary Figure S6:** Dragging columns

Here, you can see the “PivotTable Fields” on the right side of Figs S5 and S6. Drag “Measure” to Column, “Time” to Row and “Pressure” to Value.

1.6 Select the whole row data at Row 6 and Row 63.

Select the whole row data at Row 7 and Row 64, copy them to a new sheet, these are data that the Pneumatron measured every time at the 1.5 s to 30 s after pumping, and they are taken as the initial pressure ($P_i$) and final pressure ($P_f$).

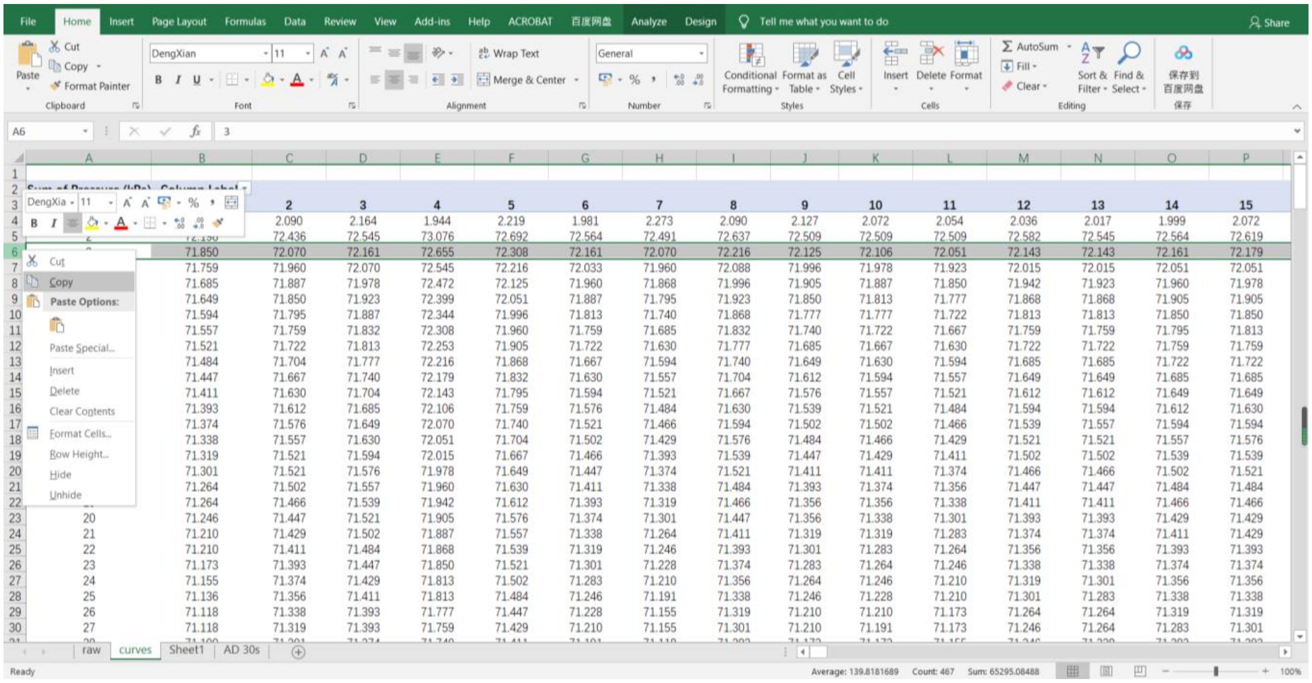

**Supplementary Figure S7**: Selecting rows

1.7 For the next steps, you may use the data_example.xlsx file in Supplementary Material 7.

**1.8** Calculate the volume of gas discharged (GD) from a branch as well as percentage of air discharged (PGD).

**Ideal gas calculations:** Ideal gas law was applied to calculate the volume of gas discharged. The moles of gas (Δn) discharged is first calculated, and it equals to:

$$\Delta n = \frac{P_i\, V_r}{R\, T} - \frac{P_f\, V_r}{R\, T} \quad (1)$$

where $V_r$ is reservoir volume (mL), which corresponds to the volume of the tubing from the pressure sensor to the connection of tube to the sample; T is the absolute temperature of gas in Kelvin, and R is the ideal gas constant (8.3144621 J mol$^{-1}$ K$^{-1}$). Convert the moles of gas discharged to volume, find the largest and smallest volume of air discharged and calculate PGD according to the equation 2:

$$PGD = 100\left(\frac{GD - GD_{min}}{GD_{max} - GD_{min}}\right) \quad (2)$$

J2 =(((I2*1000)*(E2*10^-6))/(8.3144621*293.15))-(((H2*1000)*(E2*10^-6))/(8.3144621*293.15))

| | E | F | G | H | I | J | K | L | M |
|---|---|---|---|---|---|---|---|---|---|
| 1 | Reservoir volume (mL) | Pi (kPa) | Pf (kPa) | Absolute Pi (kPa) | Absolute Pf(kPa) | GD (mol) | GD (uL) | PGD (%) | Xylem water potential (MPa) |
| 2 | 1.305 | 72.161 | 70.953 | 27.839 | 29.047 | 6.467E-07 | 16.083 | 0.0 | -0.15 |
| 3 | 1.305 | 72.161 | 70.953 | 27.839 | 29.047 | 6.467E-07 | 16.083 | 0.0 | -0.21 |
| 4 | 1.305 | 72.161 | 70.953 | 27.839 | 29.047 | 6.467E-07 | 16.083 | 0.0 | -0.27 |
| 5 | 1.305 | 72.655 | 71.264 | 27.345 | 28.736 | 7.446E-07 | 18.520 | 0.9 | -0.33 |
| 6 | 1.305 | 72.308 | 70.935 | 27.692 | 29.065 | 7.349E-07 | 18.277 | 0.8 | -0.38 |
| 7 | 1.305 | 72.161 | 70.697 | 27.839 | 29.303 | 7.838E-07 | 19.495 | 1.3 | -0.44 |
| 8 | 1.305 | 72.070 | 70.642 | 27.930 | 29.358 | 7.642E-07 | 19.008 | 1.1 | -0.50 |

K2 =(J2*8.3144621*293.15/98000)*1000*1000*1000

| | E | F | G | H | I | J | K | L | M |
|---|---|---|---|---|---|---|---|---|---|
| 1 | Reservoir volume (mL) | Pi (kPa) | Pf (kPa) | Absolute Pi (kPa) | Absolute Pf(kPa) | GD (mol) | GD (uL) | PGD (%) | Xylem water potential (MPa) |
| 2 | 1.305 | 72.161 | 70.953 | 27.839 | 29.047 | 6.467E-07 | 16.083 | 0.0 | -0.15 |
| 3 | 1.305 | 72.161 | 70.953 | 27.839 | 29.047 | 6.467E-07 | 16.083 | 0.0 | -0.21 |
| 4 | 1.305 | 72.161 | 70.953 | 27.839 | 29.047 | 6.467E-07 | 16.083 | 0.0 | -0.27 |
| 5 | 1.305 | 72.655 | 71.264 | 27.345 | 28.736 | 7.446E-07 | 18.520 | 0.9 | -0.33 |
| 6 | 1.305 | 72.308 | 70.935 | 27.692 | 29.065 | 7.349E-07 | 18.277 | 0.8 | -0.38 |
| 7 | 1.305 | 72.161 | 70.697 | 27.839 | 29.303 | 7.838E-07 | 19.495 | 1.3 | -0.44 |
| 8 | 1.305 | 72.070 | 70.642 | 27.930 | 29.358 | 7.642E-07 | 19.008 | 1.1 | -0.50 |

L2 =(100*(K2-(MIN(K:K))))/((MAX(K:K))-(MIN(K:K)))

| | E | F | G | H | I | J | K | L | M |
|---|---|---|---|---|---|---|---|---|---|
| 1 | Reservoir volume (mL) | Pi (kPa) | Pf (kPa) | Absolute Pi (kPa) | Absolute Pf(kPa) | GD (mol) | GD (uL) | PGD (%) | Xylem water potential (MPa) |
| 2 | 1.305 | 72.161 | 70.953 | 27.839 | 29.047 | 6.467E-07 | 16.083 | 0.0 | -0.15 |
| 3 | 1.305 | 72.161 | 70.953 | 27.839 | 29.047 | 6.467E-07 | 16.083 | 0.0 | -0.21 |
| 4 | 1.305 | 72.161 | 70.953 | 27.839 | 29.047 | 6.467E-07 | 16.083 | 0.0 | -0.27 |
| 5 | 1.305 | 72.655 | 71.264 | 27.345 | 28.736 | 7.446E-07 | 18.520 | 0.9 | -0.33 |
| 6 | 1.305 | 72.308 | 70.935 | 27.692 | 29.065 | 7.349E-07 | 18.277 | 0.8 | -0.38 |
| 7 | 1.305 | 72.161 | 70.697 | 27.839 | 29.303 | 7.838E-07 | 19.495 | 1.3 | -0.44 |
| 8 | 1.305 | 72.070 | 70.642 | 27.930 | 29.358 | 7.642E-07 | 19.008 | 1.1 | -0.50 |

**Supplementary Figure S8:** Volume calculations

### 1.9 Plot PGD against time/ measurements

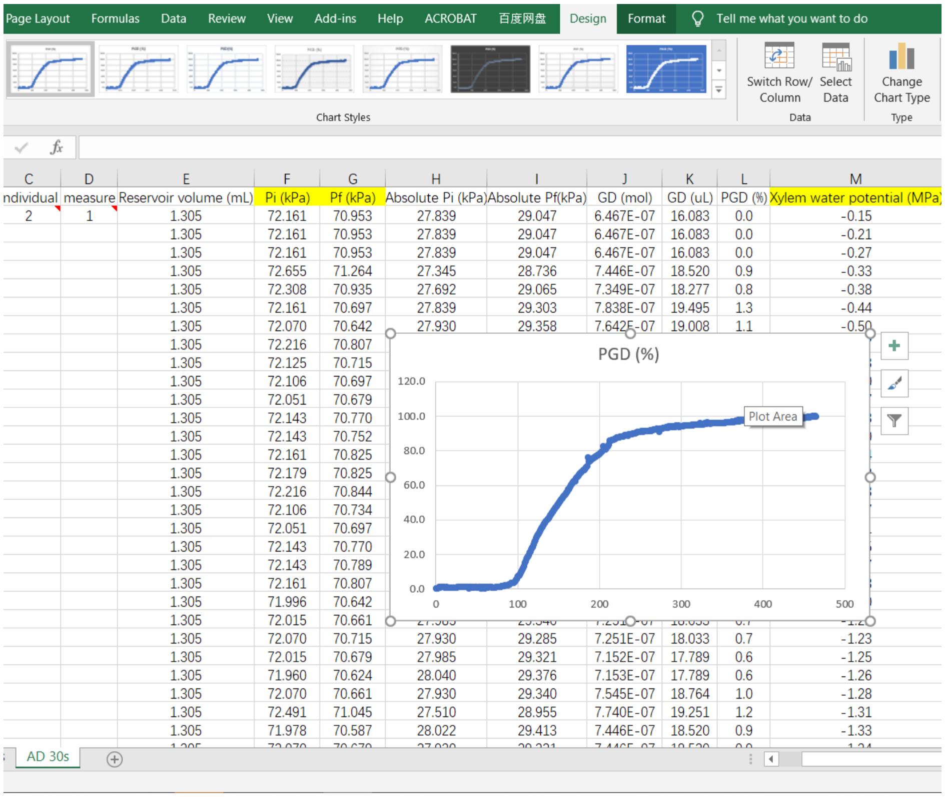

**Supplementary Figure S9:** Plotting PGD against time/ measurements

### 1.10 Get Water Potential

Get water potential results from either pressure chamber or psychrometer measurements. Plot water potential data as X-axis and the corresponding PGD results as Y-axis to get a vulnerability curve. Pale blue line indicated the sigmoidal fitting curve, and this can be fitted according the tips shown in the template.

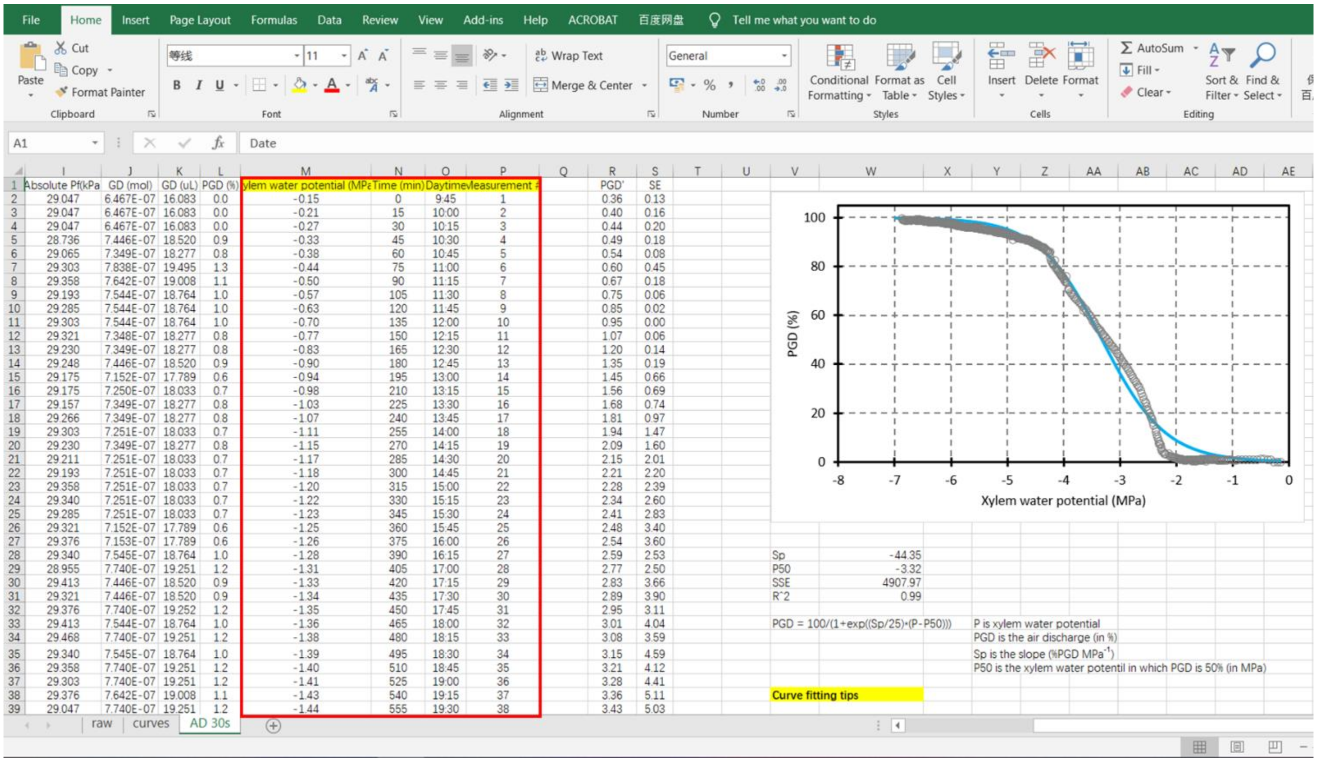

**Supplementary Figure S10:** Plotting PGD against corresponding water potential data
